# Towards solving the conundrum of plasmid mobility: networks of functional dependencies shape plasmid transfer

**DOI:** 10.1101/2022.07.04.498229

**Authors:** Manuel Ares-Arroyo, Charles Coluzzi, Eduardo P.C. Rocha

## Abstract

Plasmids are key drivers of bacterial evolution by transferring genes between cells via conjugation. Yet, half of the plasmids lack all protein coding genes for this process. We searched to solve this conundrum by identifying conjugative origins of transfer over thousands of plasmids and chromosomes of *Escherichia coli* and *Staphylococcus aureus*. We found that plasmids carrying these sequences are very abundant and have the highest densities of antimicrobial resistance genes. They are hyper-parasites that directly hijack conjugative or mobilizable elements, but not both. These functional dependencies explain the co-occurrence of each type of plasmid in cells and illuminate the evolutionary relationships between the elements. We characterized systematically the genetic traits of plasmids in relation to conjugation and alternative mechanisms of transfer, and can now propose a confident putative mechanism of transfer for ca. 90% of them. The few exceptions could be passively mobilized by other processes. We conclude there is no conundrum concerning plasmid mobility.

## Introduction

Plasmids are extra-chromosomal DNA molecules and are key drivers of horizontal gene transfer between bacteria^1^, contributing to the spread of antimicrobial resistance, virulence factors, and metabolic traits^2^. They are horizontally transmitted by several processes^3^. Some plasmids can be transferred passively, *i.e*. without dedicated genetic determinants encoded in the plasmid, by natural transformation^4^, in vesicles^5^, or by transducing bacteriophages (phages)^6^. Some plasmids are also phages, phage-plasmids (P-P), and transfer by producing their own viral particles where they package their DNA^7^. Yet, conjugation is widely regarded as the major mechanism of plasmid transfer^8^.

Conjugation involves the recognition by the relaxase (MOB) of a small DNA sequence in the plasmid, the origin of transfer (*oriT*)^9^. The relaxase cleaves the *oriT* at the *nic* site and binds covalently to the single-stranded DNA. This nucleoprotein complex, named relaxosome, interacts with a type 4 coupling protein that connects it to the mating pair formation (MPF), including a Type 4 Secretion System (T4SS) that transfers the nucleoprotein complex to another cell^10^. Once the relaxosome has been transferred, the relaxase catalyzes the DNA ligation of the plasmid in the recipient cell to produce a circular single stranded molecule that is replicated by the replication machinery of the recipient cell^9^. At the end of conjugation there is one copy of the plasmid in each cell. Some conjugative elements remain in cells as plasmids whereas others integrate the chromosome as integrative conjugative elements (ICEs)^11^. The conjugation machineries of ICEs and plasmids are very similar and have intermingled evolutionary histories^12^.

Plasmids or integrative elements encoding the three functional elements - *oriT*, relaxase and MPF - may conjugate autonomously between bacteria. They are called *conjugative*^8^. However, plasmids encoding the MPF represent only ~1/4 of all plasmids. Those lacking an MPF but encoding a relaxase and *oriT* are called *mobilizable*. In this case, the relaxase interacts with the plasmid *oriT*, and the resulting nucleoprotein complex is transported by the MPF of a conjugative element co-occurring in the donor cell. Plasmids encoding a relaxase but lacking a complete MPF are as numerous as the conjugative plasmids^8^. This means that half of all plasmids lack a relaxase and an MPF. We will refer to them as pMOBless plasmids hereinafter. Even though pMOBless lack all proteins required for conjugation, there is epidemiological evidence that some of them transfer between cells^13–15^. The mobility of pMOBless may occur by several mechanisms: (1) they may have an *oriT* and be mobilized by a relaxase and an MPF encoded *in-trans* by a conjugative plasmid^16^; (2) they may interact with a relaxase of a mobilizable plasmid, and the nucleoprotein complex further interacts with an MPF of a third plasmid^17^; (3) or they may transfer using other mechanisms, *e.g*. conjugation through a rolling circle replication protein^18^, co-integration with a conjugative plasmid^19^, or the alternative transfer mechanisms mentioned above. Similar mechanisms could be used by integrative elements lacking a complete MPF, commonly named integrative mobilizable elements (IMEs)^20^.

The observation over a decade ago that slightly more than half of all plasmids lack genes for relaxases was paradoxical, because genetic mobility is thought to be necessary for plasmid maintenance in populations^21,22^. Of note, some pMOBless with an *oriT* (pOriT hereinafter) were shown to be mobilized by a conjugative plasmid decades ago^17^. Yet, the few available sequences of *oriT* have precluded systematic identification of these plasmids. Recently, pioneering studies on *Staphylococcus aureus*, a species that has unusually few conjugative plasmids and few types of *oriT*, showed that 50% of the pMOBless can be mobilized since they carry *oriTs* similar to those of pWBG749^23^ or pSK41^24^. Subsequent studies with three additional *oriT*s, suggested that *oriT*-based mobilization is common in this species^25,26^. If this is true for other species, including those with numerous conjugative plasmids, is not known. Unfortunately, most *oriTs* remain unknown, precluding their systematic study across bacteria. Here, we focused on *S. aureus*, for which plasmid diversity is low and well-characterized and *Escherichia coli*, the best described species of bacteria and one with numerous well-known plasmid families^27^. These two species are of particular importance because they are responsible for the greatest number of deaths associated to antimicrobial resistance in the world^28^, a trait that is spread by plasmids^29^. We first complement previous studies and test if ICEs could be involved in the mobilization of pOriTs in *S. aureus*. We also test if the same approach can be extended to *E. coli*. The confirmation that we can identify homologs of experimentally verified *oriT*s in the plasmids of these species paved the way to answer some outstanding questions. We don’t know how these plasmids contribute to the spread of functions across bacteria. We don’t know the functional dependencies associated with pOriTs, *i.e*. if they tend to be associated with one single conjugative plasmid or if they often require a third plasmid encoding a relaxase. We don’t know how these plasmids arose in natural history. We also ignore how the existence of pOriTs affects the patterns of co-occurrence of plasmids in cells. Finally, we would like to know how many plasmids remain without a hypothetical mechanism of transfer once pOriT plasmids and phage-plasmids are accounted for. By tackling these questions, this study contributes to unravel the mechanisms allowing plasmid mobility, while giving new insights into the mobility and evolution of *oriT*-bearing plasmids.

## Results

### *E. coli and S. aureus* have distinct plasmid repertoires

We analyzed the complete genomes available in RefSeq of *E. coli* (n=1,585) and *S. aureus* (n=581) to characterize the size and diversity of their plasmids. *E. coli* isolates carry almost three times more plasmids per genome than *S. aureus* isolates (t_(2068.9)_=20.65; p<2.2e-16) (Fig 1A). Moreover, *E. coli* plasmids tend to be larger (Kolmogorov-Smirnov test, D=0.586, p<2.2e-16) (Fig 1B) and with a higher GC% than *S. aureus* plasmids (t_(1074.7)_=191.23, p<2.2e-16) (Fig S1). They are also more diverse in terms of gene repertoires. *E. coli* plasmids encode on average four times more gene families than those of *S. aureus* (t_(2817.9)_=43.129, p<2.2e-16) (Fig S1). The plasmid pangenome of *E. coli* (11,530 gene families) is much larger than that of *S. aureus* (ca. 1,000), a trend that could be confirmed when comparing similar sampling sizes (455 plasmids) (Fig 1C). Overall, plasmids contribute with many genes to the species pangenomes. This is particularly striking in *E. coli*, where the plasmid pangenome is more than double the average size of a strain genome^30^.

**Figure 1.**
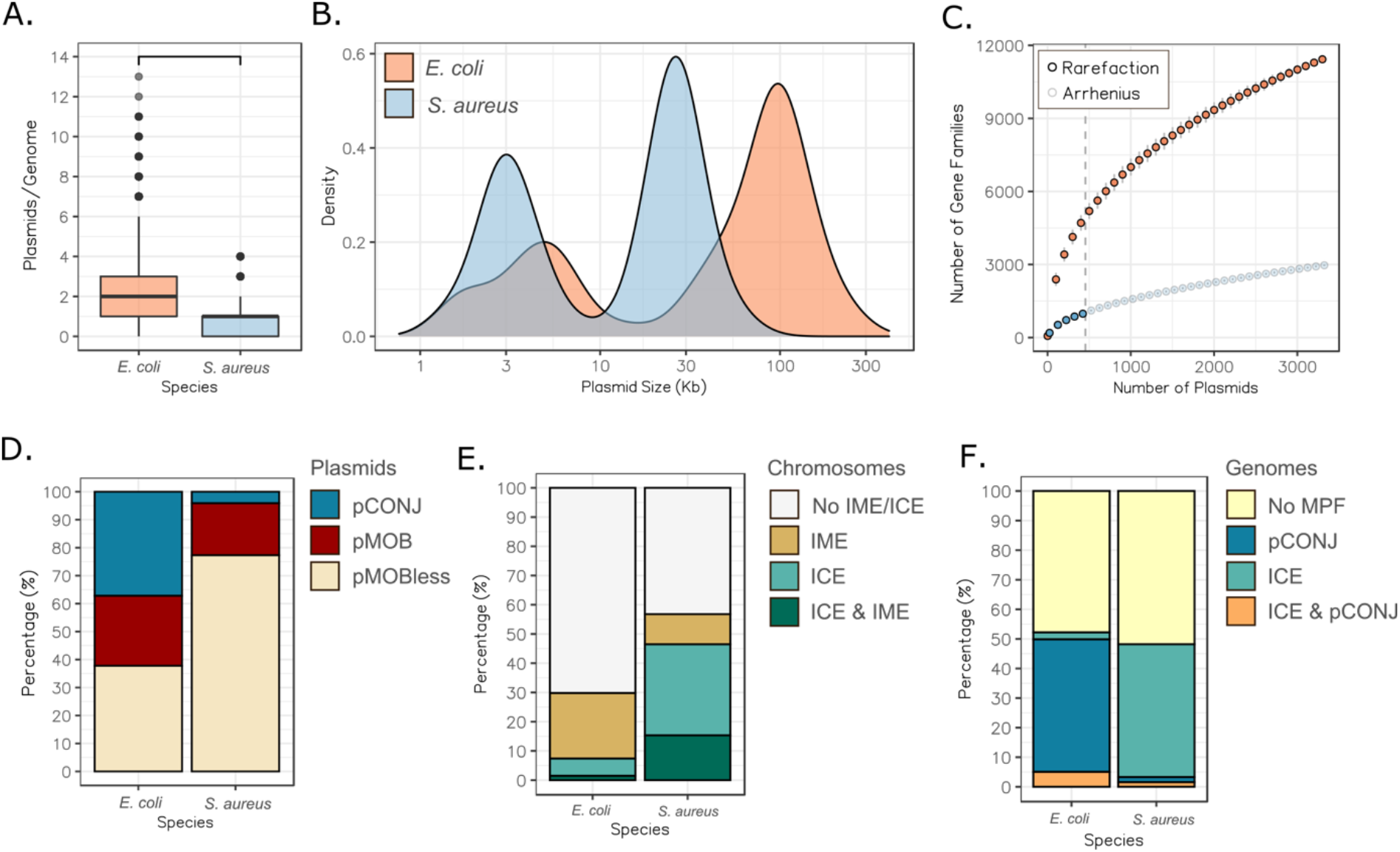
Comparison of *E. coli* and *S. aureus* plasmids. **A.** Number of plasmids per genome. The horizontal bar over the plot denotes statistically significant difference (t_(2068.9)_=20.65; p<0.0001). **B.** Plasmid size distribution. The curves were drawn using a kernel density estimate. **C.** Plasmid pangenome of *E. coli* and *S. aureus* attending to the number of plasmids sampled. The vertical dashed grey line at x=455 represents the number of plasmids from which *S. aureus* pangenome is inferred following an Arrhenius model. **D.** Percentage of each mobility category (conjugative, pCONJ; mobilizable, pMOB; presumably non-transmissible, pMOBless) among the plasmid repertoire of both species. **E.** Percentage of the chromosomes with at least one ICE (complete MPF and relaxase), IME (relaxase without a complete MPF), both ICE and IME, or no conjugative elements. **F**. Fraction of genomes with a complete MPF, either in plasmids (pCONJ), in the chromosome (ICE) or in both.

We characterized the plasmids in terms of the protein coding genes involved in conjugation: pCONJ encode an MPF and a relaxase, pMOB encode a relaxase, and pMOBless lack a relaxase. In *E. coli* ~35% of the plasmids are pCONJ, ~25% pMOB, and ~40% pMOBless (Fig 1D). These values are close to previously published ones across Bacteria^8^. In contrast, only 4% of the *S. aureus* plasmids were classed as pCONJ, 18% as pMOB, and 77% as pMOBless. Hence, *S. aureus* seems a more atypical bacteria, where conjugative plasmids are rare. We then tested the hypothesis that ICEs could compensate for the paucity of conjugative plasmids in the species. We searched the chromosomes for loci associated with ICEs (encoding MPF and relaxase) and IMEs (encoding a relaxase), and found that 46% of the chromosomes of *S. aureus* encode MPF systems (Fig 1E). In contrast, conjugative systems were identified in only ~7% of *E. coli* chromosomes. Interestingly, many genomes in both species have either conjugative plasmids or ICEs, but rarely both. The integration of these analyses provides a more nuanced view of the differences between the species in terms of the fraction of genomes containing a conjugative element: ~52% of *E. coli* and ~47% of *S. aureus* (Fig 1F). While the precise delimitation of ICEs and IMEs is difficult and precludes systematic comparisons between elements in terms of gene content, these results suggest that the existence of ICEs could explain the mobility of some pMOBless, especially in *S. aureus*. In summary, the two species show different patterns in terms of the mobility of plasmids and integrative elements, but both still contain many plasmids lacking relaxases.

### *oriTs* are frequent in plasmids of *E. coli* and *S. aureus*

To unveil the mechanisms of mobilization of the many plasmids lacking a relaxase, we searched for *oriTs*. To do so, we collected 51 *oriT* from the *‘oriT* database’^31^ and added 40 new ones from the literature (Table S3). Most of these 91 experimentally validated *oriTs* (mean size ~131 bp) were originally identified and verified in plasmids of γ-Proteobacteria (n=44) and Bacilli (n=22) (Fig S2). We used it to search for origins of transfer in the 1,585 *E. coli* and 581 *S. aureus* genomes by sequence similarity (see Methods). We identified 2,831 putative *oriT*s in 2,626 plasmids, almost the totality of which locate in intergenic regions (Fig S3). Even if *E. coli* has more diverse plasmids and more types of *oriTs* (n=37) than *S. aureus* (n=7), *oriTs* were found at similar frequencies in the plasmids of the two species (ca. 70%) (Fig 2A). We also identified 336 *oriT*s in 282 chromosomes. These chromosomal *oriT* were much more abundant in *S. aureus* (25% of the genomes) than in *E. coli* (9%), in line with the higher frequency of ICEs in the former (Fig 2A). Although many *oriT*s were identified in both types of replicons, a given family tends to be present either in plasmids or in chromosomes (Fig 2B). To note, none of the *oriT*s was identified in both species.

**Figure 2.**
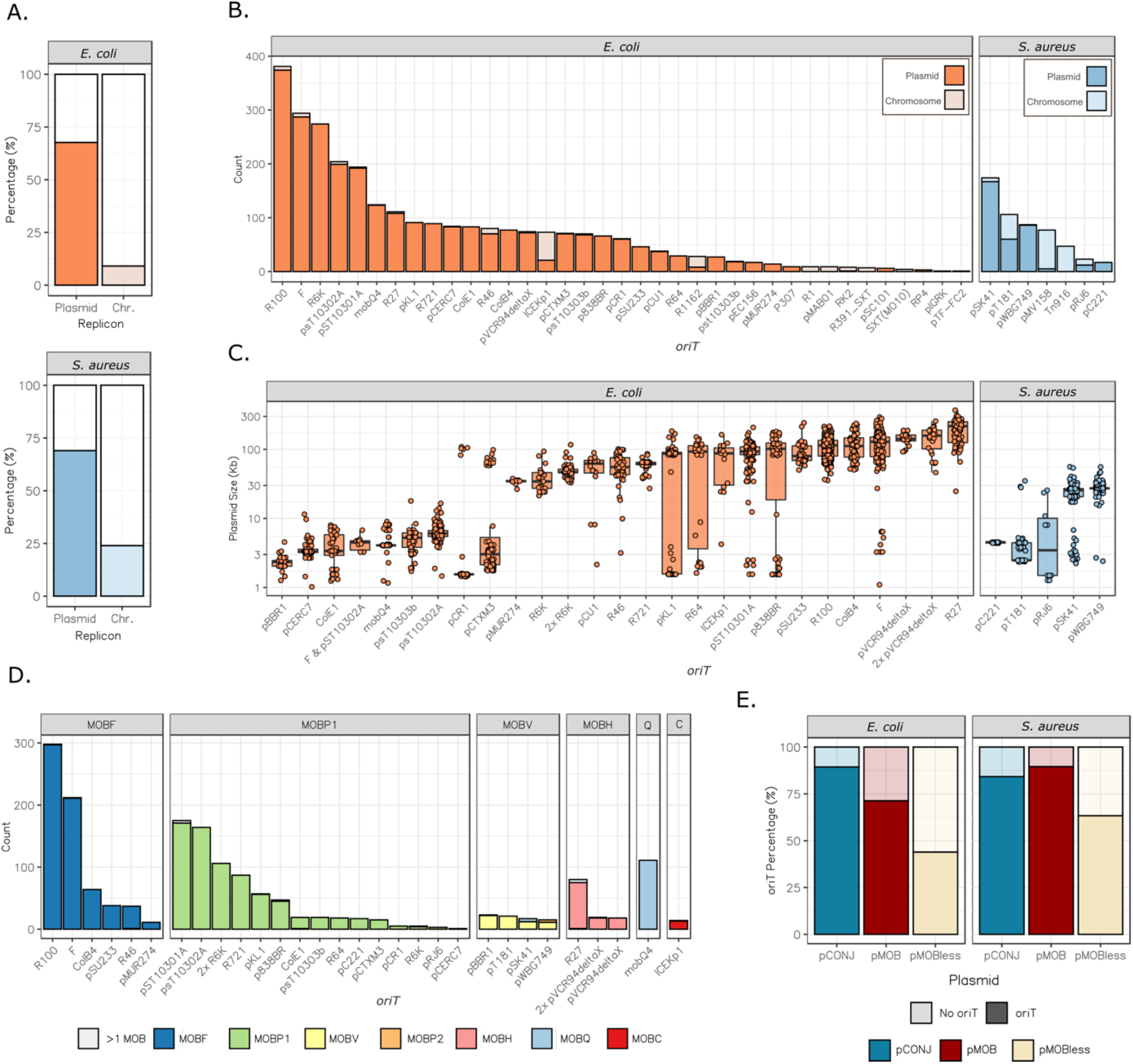
Identification of *oriTs*.**A.** Proportion of plasmids and chromosomes with at least one *oriT* in *E. coli* (top) and *S. aureus* (down). **B.** Counts of *oriTs* in the genomes of *E. coli* (left) and *S. aureus* (right). **C.** Size of plasmids containing an *oriT* (or a combination of *oriT*s) present in at least 10 plasmids. **D.** MOB families associated to the *oriTs* in (C.). **E.** Percentage of plasmids in which at least one *oriT* was identified, classed by mobility type.

Most *oriT*-encoding plasmids have just one *oriT* (~88%*E. coli*, ~85%*S. aureus*), although a few can have up to 5 (Fig S3). Expectedly, plasmids showing multiple *oriTs* tend to encode multiple relaxases (r(3868)=0.32, p<2.2e-16) (Fig S3). To study the plasmid size and the co-occurrence of *oriTs* and relaxases, we retrieved the families of *oriT*s identified in more than 10 plasmids. The *oriT*s of a given family are usually associated with plasmids of a specific size range, *i. e*., they tend to be associated to either small or large plasmids (Fig 2C). Yet, in a few cases, the families associated with large plasmids also include a few much smaller ones. Finally, the *oriT*s of a given family tend to be in plasmids with the same class of relaxases (Fig 2D). All things considered, the identification of *oriTs* in most plasmids, usually in a single copy, the strict association between the *oriT* and the MOB, and their identification in plasmids of homogeneous size, suggest that most *oriTs* we identified are true positives.

### *oriT*-MOBless plasmids are abundant and usual carriers of antimicrobial resistance genes

We identified at least one *oriT* in more than 80% of pCONJ and pMOB (Fig 2E). Hence, the *oriTs* in our collection have homologous sequences in a very large fraction of the *oriTs* used by the conjugative plasmids of these species. Importantly, we found an *oriT* in 790 pMOBless. Hereinafter, we will refer to these *oriT*-carrying pMOBless as pOriT. pOriTs constitute 65% of *S. aureus* plasmids lacking relaxases and more than 40% of those of *E. coli*. These results are subject to caution. We cannot ascertain the functionality of all these *oriT*, even if they are homologous to experimentally verified sequences. More importantly, our analysis may still be missing *oriT*s, since even a few pCONJ lack an identifiable *oriT*. Despite these limitations, most plasmids have one and only one identifiable *oriT*, suggesting that we have identified most of them. If so, around half of the plasmids lacking relaxases are mobilizable by conjugation.

Due to the importance of *E. coli* and *S. aureus* as multidrug resistant pathogens^28^, we enquired on the role of their different plasmids in the spread of antimicrobial resistance genes (ARG). It has previously been found that conjugative plasmids tend to carry more ARGs than the other plasmids^29^. This is the case of pCONJ in *E. coli* (~64% of the genes) but not in *S. aureus*, where pOriTs carry most of these genes (~76%) (Fig 3A). Furthermore, the number of ARGs per kilobase is highest in pOriT in both species (Fig 3B). Interestingly, the plasmids with fewer ARGs, and lowest density, are those lacking both a relaxase and an *oriT* (presumably non-transmissible, pNT). These results show that plasmids lacking relaxases can be split in two categories, where those with an *oriT* have an important role in the spread of antibiotic resistance.

**Figure 3.**
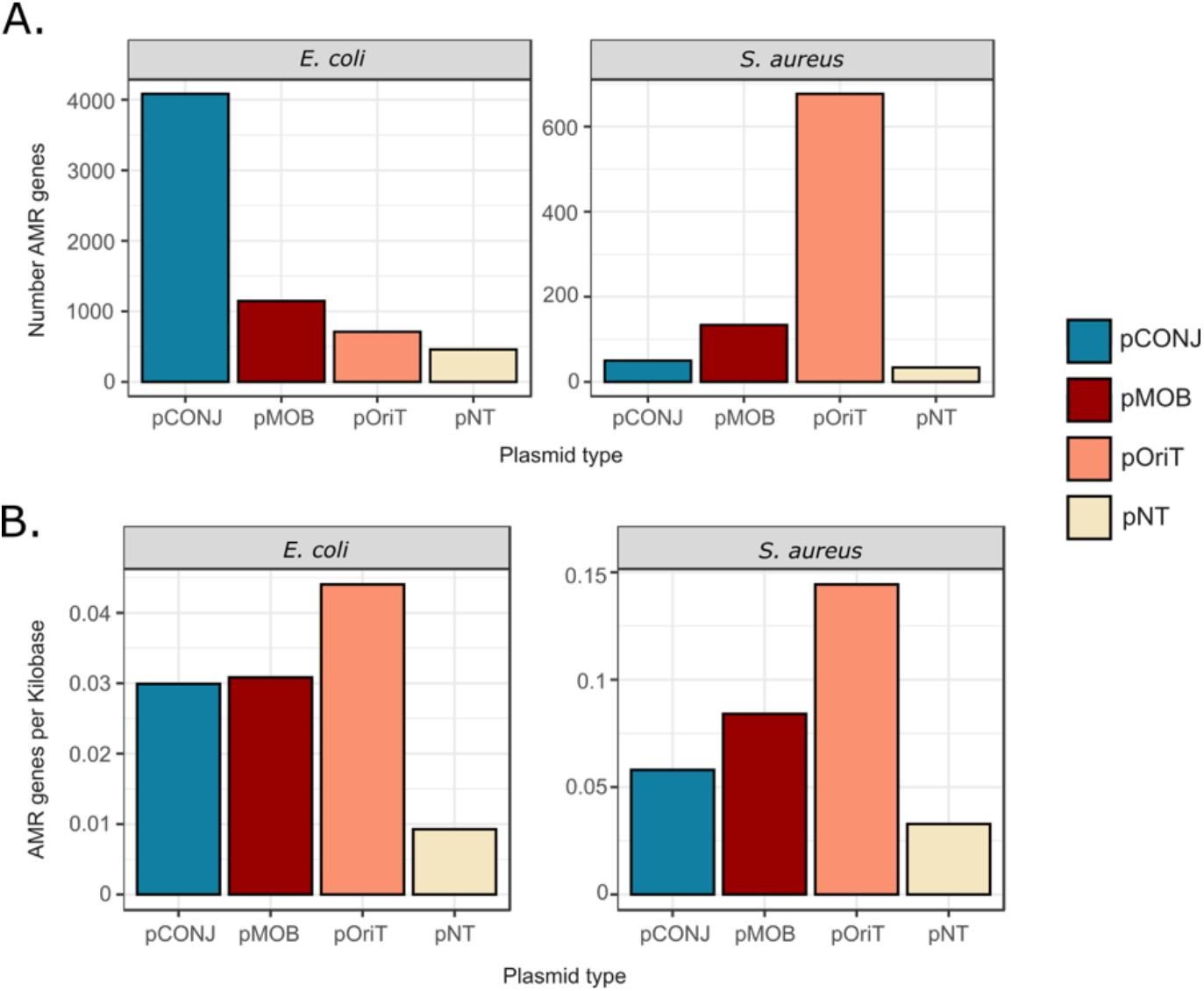
Plasmid types and antimicrobial resistance (AMR). **A.** Number of AMR genes encoded in each plasmid type. **B.** Density of AMR genes (genes per kilobase) according to the plasmid type.

### pOriTs exploit either conjugative or mobilizable plasmids

The identification of homologous *oriT*s allows to test functional dependencies between plasmids. We have previously proposed that relaxases of pMOB evolve to interact with multiple types of MPF encoded in pCONJ, whereas those of pCONJ co-evolve with the MPF to optimize their mutual interaction^32,33^. These differences might require the presence of different *oriT*s in pMOB and pCONJ, as previously suggested^26^. In our dataset, many families of *oriT*s are present in either pCONJ or pMOB, but few are present in both (Fig 4A). The exceptions tend to correspond to “pCONJ-like *oriTs*” (*oriTs* typical of pCONJ) that were found in large pMOB plasmids. We hypothesized that these might be decayed conjugative plasmids (pdCONJ)^34^. These elements have some MPF genes, but not enough to be functional, and seem to have been recently derived from pCONJ by gene deletion^34^. Hence, we split the pMOB into those encoding at least two MPF genes (pdCONJ) and the others. The pdCONJ are indeed 80% of the mobilizable plasmids with pCONJ-like *oriTs*. In contrast, pdCONJ do not have “pMOB-like *oriTs*” (*oriTs* typical of pMOB) (Fig 4A). After this analysis, only three *oriTs* remained in a significant fraction of both pCONJ and pMOB (excluding pdCONJ): *oriT*_pKL1_, *oriT*_pWBG749_, and *oriT*_pSK41_. We then enquired on the possibility that ICEs or IMEs show similar trends. Since we ignore the limits of these elements, we cannot properly assign them an *oriT*. Yet, we can analyze if certain *oriT*s are present in chromosomes encoding an ICE or/and an IME. Our results showed that indeed, *oriT*s tend to be associated with either ICEs or IMEs (Fig S4). We conclude that conjugative and mobilizable elements tend to use different *oriT*s.

**Figure 4.**
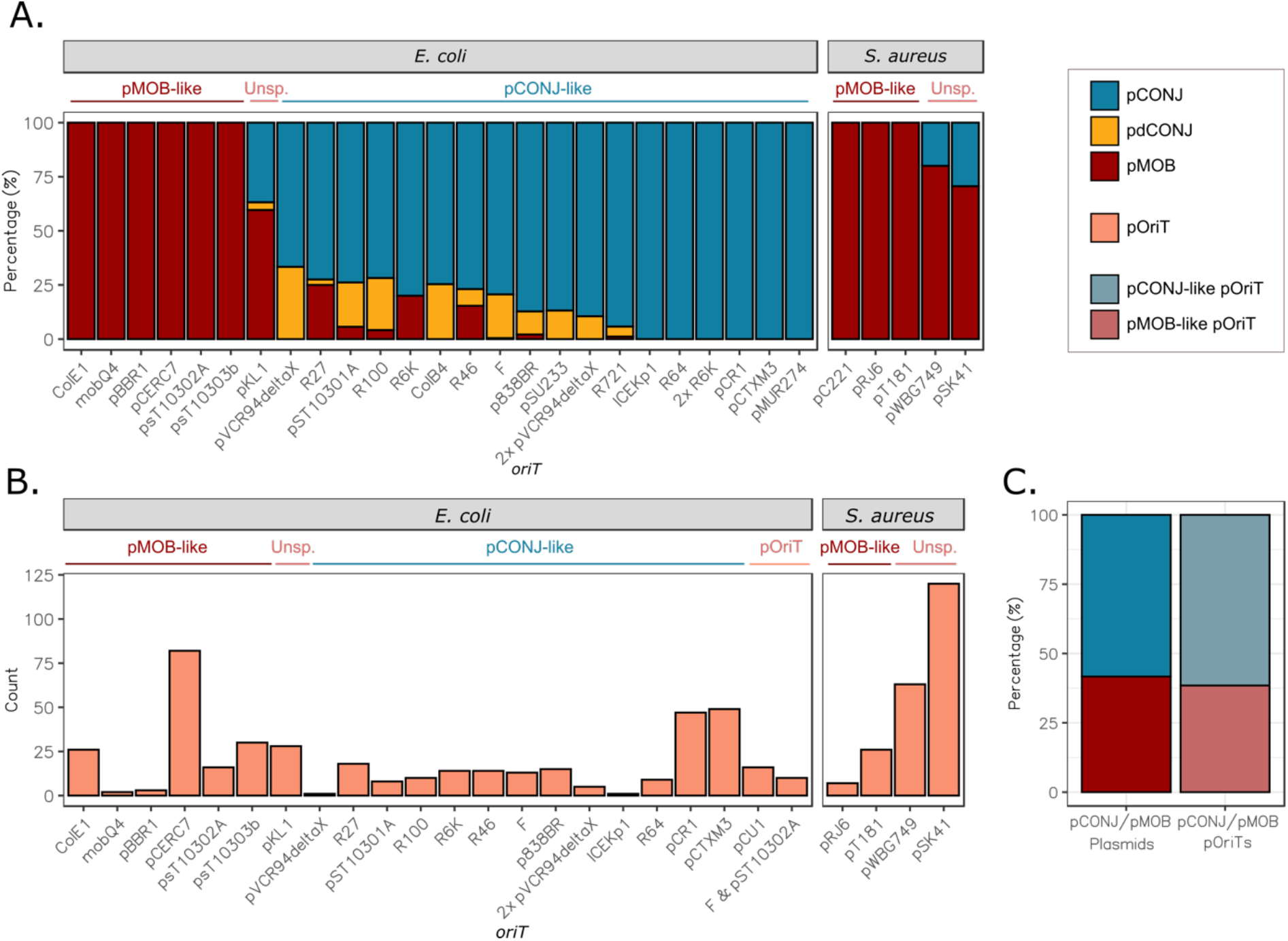
**A.** Proportion of plasmid types having a given *oriT* or a combination of *oriTs* (for those occurring in more than 10 plasmids). **B.** Number of pOriTs (*oriT*-encoding MOBless plasmids) found for each *oriT*. pCONJ-like: conjugative *oriT*, identified mostly (>75%) in conjugative plasmids; pMOB-like: mobilizable *oriT*, identified mostly (>75%) in mobilizable plasmids; Unsp.: *oriT* identified in many conjugative and mobilizable replicons; pOriT: *oriT*s identified only in pOriTs. The color indicates the plasmid mobility, being the legend at the top right of the figure. **C.** Ratio of pCONJ/pMOB plasmids compared to the ratio of pOriTs with pCONJ-like and pMOB-like *oriT*s in *E. coli*.

A plasmid encoding only an *oriT* may either use the relaxase and MPF of a conjugative plasmid (if carrying a pCONJ-like *oriT*), or the relaxase of a mobilizable plasmid which in turn must use an MPF of a conjugative one (if carrying a pMOB-like *oriT*). In the first case, the pOriT could be regarded as a parasite of the conjugative plasmid, if its activity affects the fitness of the latter, whereas in the second case it is a hyper-parasite (a parasite of a parasite). One could expect that the most efficient strategy for a pOriT would be to take advantage of a unique plasmid rather than relying on the interplay between two other elements. However, since pMOB are often able to interact with multiple pCONJ, a pMOB-like *oriT* might allow a pOriT to have a higher chance of transfer under certain circumstances. Since the *oriT*s of pOriTs are homologous to those of conjugative or mobilizable elements (Fig 4B), we could infer the relations of dependence between pOriT and the other plasmids. We focused on *E. coli* plasmids for this particular analysis because they have a much wider diversity of *oriT*s for both pMOB and pCONJ. Interestingly, the frequency of pOriTs in *E. coli* with a pCONJ-like *oriT* (~56%) or a pMOB-like one (35%) is very close to the relative frequency of each of these types of plasmids in the species (Figure 4C). Hence, the relative frequency of each type of pOriT matches the relative frequency of the hijacked plasmids.

### pOriT may originate from both conjugative and mobilizable plasmids

Given the large number of pOriTs, we enquired on their evolutionary origin. It was recently suggested that pMOBless may have derived from conjugative or mobilizable plasmids by gene deletion^34^. Since pOriTs have either a pCONJ-like or a pMOB-like *oriT*, we thought they might have emerged by gene deletion in ancestral pCONJ or pMOB while maintaining the *oriT*. To evaluate this hypothesis, we grouped the 3,869 plasmids into Plasmid Taxonomic Units (PTUs)^27^ and analyzed their mobility and *oriT*. Most plasmids in a PTU have the same type of mobility, reflecting the short evolutionary distances between plasmids in the same PTU. But even when they do not, they tend to have *oriT*s of the same family (Fig S5), suggesting that *oriT* family is more conserved than the mobility type.

To test the possibility that some pOriTs originated from conjugative plasmids, we selected two PTUs and explored the relation between the pOriTs and pCONJ within a PTU. We analyzed the PTU-F_e_ (IncF/MOB_F_/MPF_F_) (Fig 5) and the PTU-C (IncA/C2/MOB_H_/MPF_F_) (Fig S6). Most of the plasmids in these PTUs are pCONJ with a pCONJ-like *oriT* (*oriT*_F_ and *oriT*_pVCR94deltaX_, respectively). Yet, both include a few other types of plasmids (*e.g*. pMOB, pOriT) that tend to be smaller than their pCONJ counterparts (PTU-Fe: F_(481)_=8.808, p=7.21e-07; PTU-C: F_(37)_=35.69, p=2.32e-09) while encoding the usual *oriT* of their PTU (Fig 5, Fig S6). This supports the idea that these replicons derived from conjugative plasmids by gene deletion. To further test this idea, we analyzed pairs of pCONJ/pOriT within the PTUs having similar gene repertoires (wGRR>0.75, see Methods). This analysis suggests that these pOriTs were generated by staggered degradation of the MPF system in pCONJ (Fig 5, Fig S6). Crucially, the derived replicons are likely to be able of *in-trans* conjugation because of the maintenance of their ancestral *oriT*.

**Figure 5.**
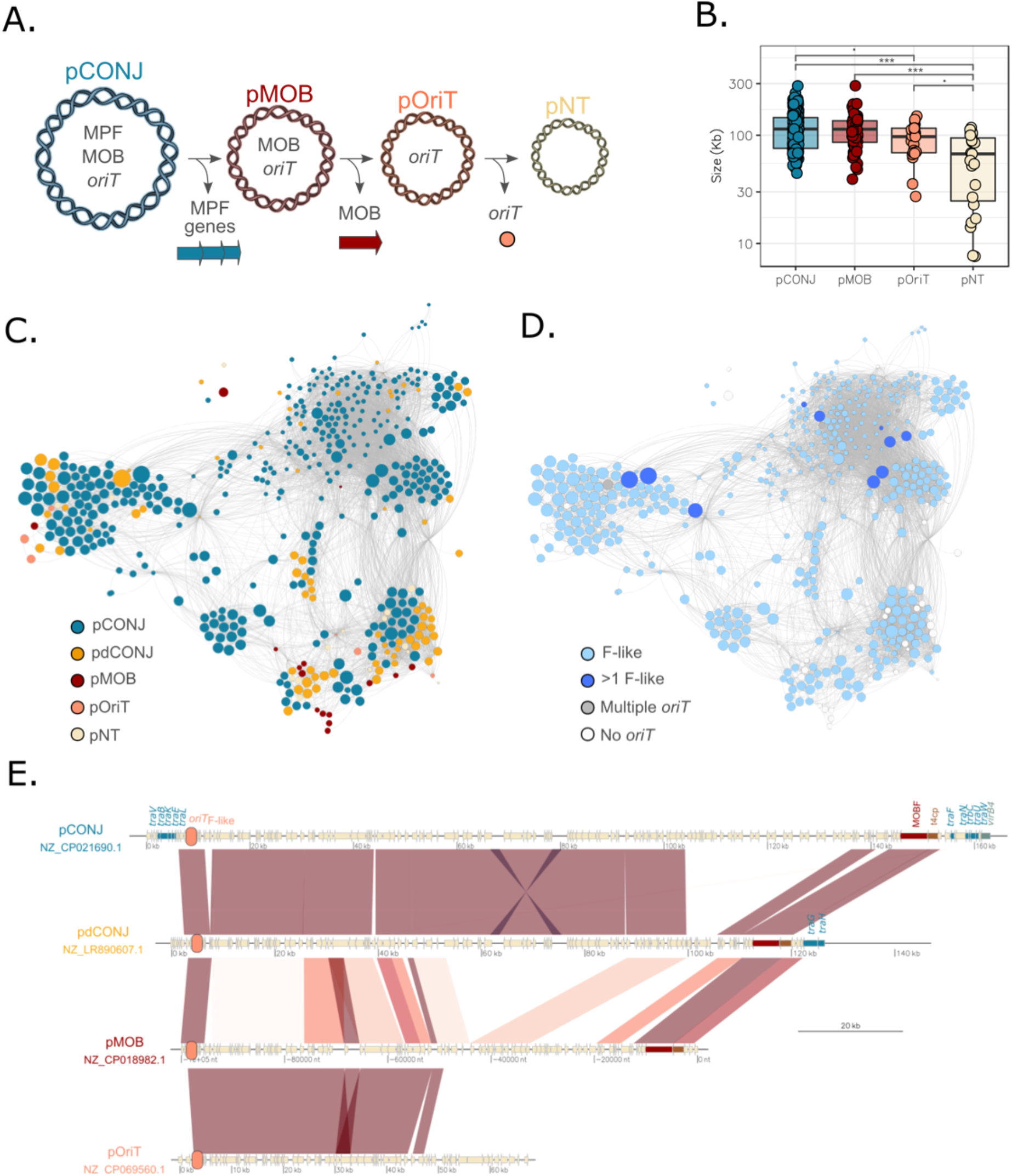
Evolution of pCONJ-like pOriTs. **A.** Proposed evolutionary hypothesis for the origin of pCONJ-like pOriTs. **B.** Plasmid size of the PTU-Fe according to their mobility. The horizontal bars over the plot denote statistically significant difference (pairwise t-tests): ***(p<0.001), **(p<0.01), *(p<0.05), ·(p<0.1). **C.** and **D.** Graphs showing the PTU-Fe. Nodes represent the plasmids and edges connect plasmid pairs with wGRR>0.75. The colors of the nodes represent the plasmid mobility (**C**) and the *oriT* (**D**). **E.** Plasmid alignments of a pCONJ, pdCONJ, pMOB and pOriT from the PTU-Fe. Conjugative genes are indicated as blue arrows, the relaxase in red, coupling protein in brown, *virB4* in green, and the *oriT* as an orange circle.

We then selected two PTUs with a majority of pMOB (E1, E22) and analyzed them as above (Fig 6, Fig S7). Both include ColE1-like plasmids (ColRNAI/Col440I), associated to the MOB_P_ and the pMOB-like family *oriT*_ColE1-like_. As before, these PTUs include other types of plasmids, notably pOriTs and pNTs. The latter tend to be smaller (PTU-E1: F_(200)_=90.33, p=<2e-16; PTU-E22: F_(35)_=827.18, p=7.53e-08), again suggesting that they arose by deletion of the relaxases in ancestral pMOBs. As expected, most of the closely related pMOB/pOriT pairs have homologous *oriT*s, and their alignments further suggest that small pOriTs arise by the loss of the relaxase in pMOB plasmids (Fig 6, Fig S7). Interestingly, we identified a change of the *oriT* from one to another family in a subgroup of plasmids of the PTU-E1 (Fig 6). This subgroup of plasmids have the *oriT*_pCERC7_, an origin of transfer related to the pCONJ-like *oriT*_R64_^35^. This finding suggests that through recombination events, a family of pMOBless with pMOB-like *oriTs* can acquire an *oriT* typical of conjugative plasmids. Overall, these results show at the micro-evolutionary scale how pOriTs can derive by gene deletion from other types of plasmids.

**Figure 6.**
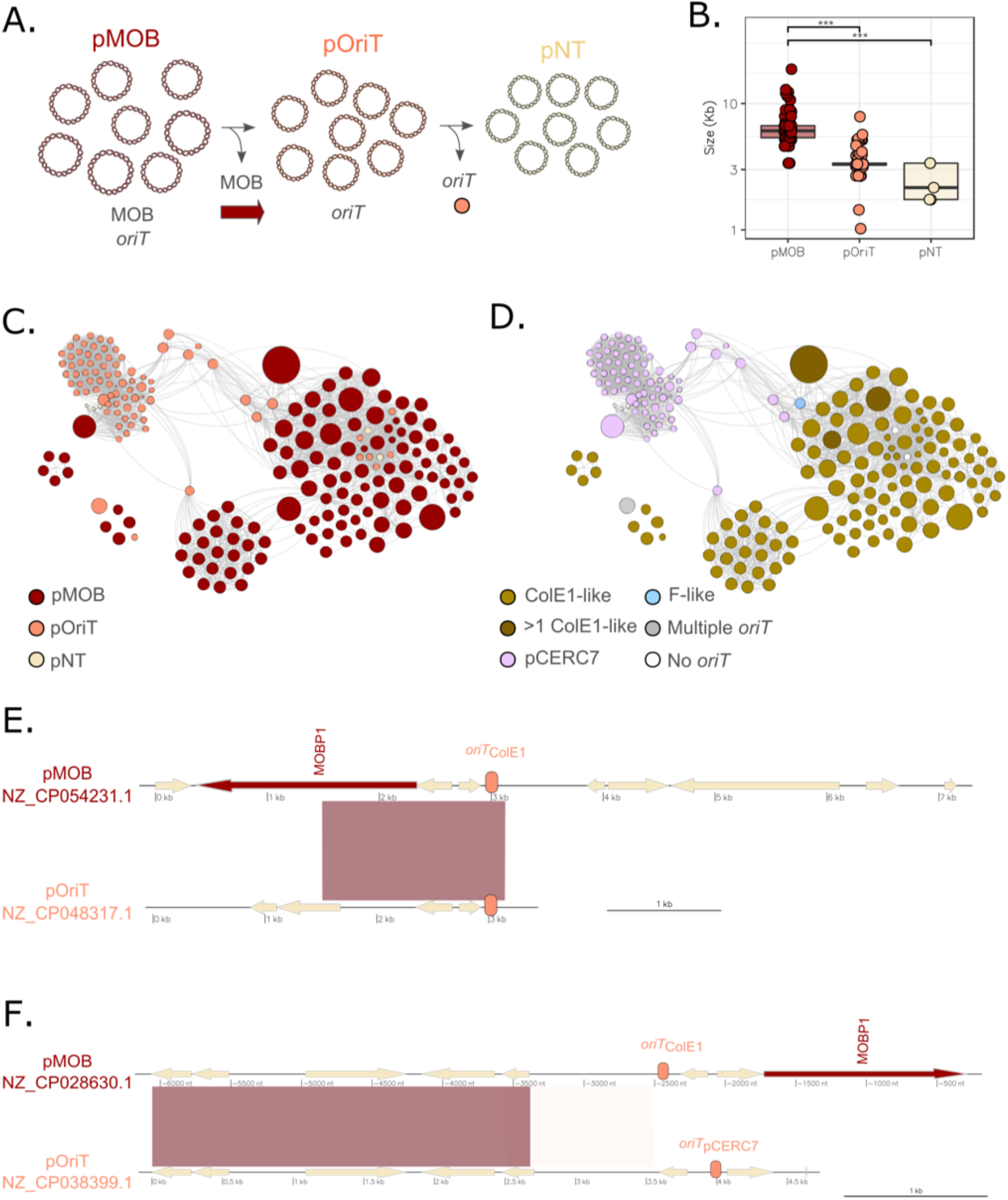
Evolution of mobilizable-like pOriTs. **A.** Proposed evolutionary hypothesis for the origin of mobilizable-like pOriTs. **B.** Plasmid size of the PTU-E1 according to their mobility. The horizontal bars over the plot denote statistically significant difference (pairwise t-tests): ***(p<0.001), **(p<0.01), *(p<0.05), ·(p<0.1). **C.** and **D.** Graphs showing the PTU-E1. Nodes represent the plasmids and edges connect plasmid pairs with wGRR>0.75. The colors of the nodes represent the plasmid mobility (**C**) and the *oriT* (**D**). **E.** and **F.** Plasmid alignments of a pMOB, and pOriT from the PTU-E1. The relaxase is indicated as a red arrow, and the *oriT* as an orange circle.

### Most plasmids may be mobilized by known mechanisms of transfer

Our results suggest that ~80% of *E. coli* and >70%*S. aureus* plasmids use an *oriT* to transfer by conjugation. To this, one may add other genetic elements that spur plasmid transfer (Fig 7A). Notably, some rolling-circle replication proteins (RC-Rep) act as replicative relaxases^36^. They interact with the MPF system of a conjugative element and trigger plasmid conjugation in an *oriT*-independent manner^37^. We searched for these proteins to test if this alternative pathway could be involved in the mobilization of plasmids lacking *oriT* and classical relaxases. We identified 225 homologs of RC-Rep proteins in 208 plasmids. These plasmids are frequent in *S. aureus* (~30%), but rare in *E. coli* (1.9%). As expected, there is an overrepresentation of RC-Rep in *non-oriT* pMOBless (χ^2^_(4)_=103.12, p<2.2e-16) (Fig S8). The unexpected abundance of RC-Rep in plasmids lacking an *oriT* suggests that such proteins could mediate the mobility of many plasmids in *S. aureus*.

**Figure 7.**
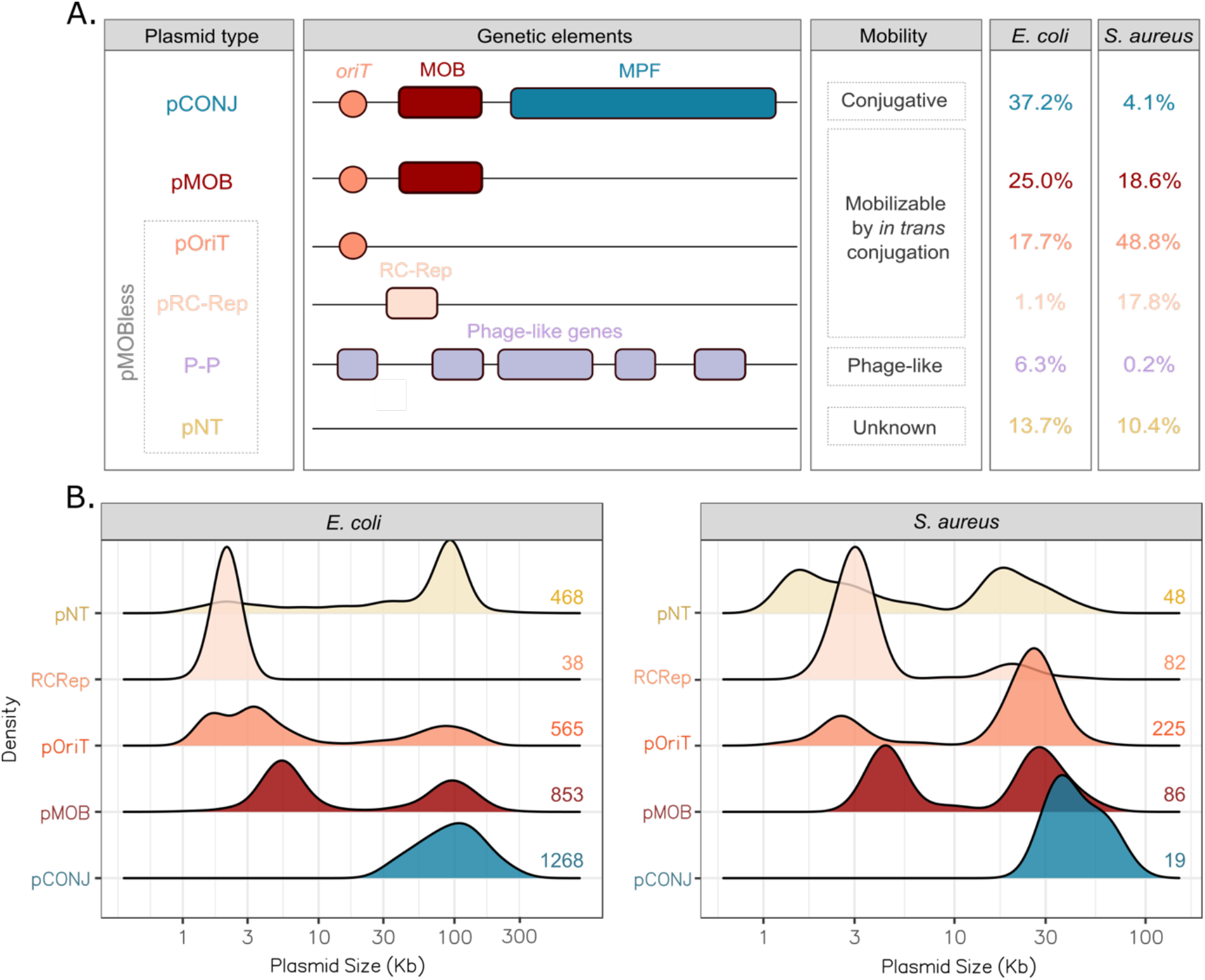
Classification of plasmid mobility. **A.** Representation of plasmids in function of their category, genetic composition, and mechanism of mobility. The frequency (%) of each plasmid type in *E. coli* and *S. aureus*,respectively, is shown at the right columns of the figure. **B.** Plasmid size attending to the mobility. The curves were drawn using a scaled kernel density to simplify the representation (sample sizes at the right of each row). The size distribution of P-Ps is shown in the Sup Fig 9.

Some plasmids can be transferred within viral particles. The propensity of a plasmid to be transduced cannot be predicted from its sequence. But ca. 6% of the plasmids are also phages (phage-plasmids, P-Ps)^7^, and encode viral particles, virion assembly packaging, and cell lysis. We identified 222 P-Ps in *E. coli* and 1 in *S. aureus*, which is consistent with the reported uneven distribution of P-Ps across bacteria^7^. P-Ps correspond to a third of the pMOBless without *oriT* in *E. coli* (n=216/702). In agreement with the idea that P-Ps provide an alternative mechanism of plasmid transfer, only six P-Ps encode conjugation-related elements (Fig S9). The latter are much larger (~175 kb) than the remaining P-Ps (~90 kb), and might be the result of co-integration events or assembly artifacts (Fig S9).

At the end of these analyses, we could assign a putative mechanism of mobility for most plasmids in each species. In *E. coli*, 80% of the plasmids were classed as conjugative or mobilizable by conjugation, and ~7% as P-Ps. In *S. aureus*, 90% were classed as conjugative or mobilizable by some type of conjugation and only 1 is a P-P. Hence, when one accounts for MPF, relaxases, RC-Rep, *oriT*, and P-Ps, few plasmids lack a hypothetical mechanism of transfer, *i. e*. few remain putatively non-transmissible (pNT) (Fig 7A): 13.7% in *E. coli* and 10.4% is *S. aureus*. We enquired on the possible mechanisms of mobility of the remaining plasmids. Around 50% of the *E. coli* pNTs are related to the large plasmid pO157 (PTU-E5) (Fig S10). These are well-known non-transmissible plasmids that have disseminated in *E. coli* O157:H7^39^. The mechanisms of mobility of the few remaining plasmids (if any) remains unknown.

The distribution of the size of plasmids is bi-modal and associated with their type of mobility (Fig 7B). The mode associated with the largest plasmids is characteristic of pCONJ, but also found among certain pMOB and pOriT in both species. For the latter, we observed a shift of the peak to lower values of plasmid size. Similarly, the mode of the smaller plasmids is characteristically associated with pMOB, but is also found among pRCR and pOriT, with a shift of the peak to lower values of plasmid size. These small downwards shifts observed among pOriT and other plasmids are consistent with our hypothesis that they often originate from pCONJ or pMOB by gene deletion (Fig S11). The patterns for pNT are less clear. In *E. coli* they are shaped by the many large pO157-like plasmids, whereas in *S. aureus* they seem to follow the trends of pOriT, suggesting that maybe some *oriT* remain to be uncovered in the species.

### Mobilization explains patterns of plasmid co-existence

The dependence of certain plasmids, *e.g*. pOriT, on others, notably pCONJ, for conjugative transfer means that the type of mobility of plasmids may affect the patterns of their co-occurrence in cells. We can now test this hypothesis by analyzing which plasmids tend to co-occur with others. The number of plasmids per genome is much more variable (and on average higher) in *E. coli* than in *S. aureus*. Hence, we concentrated on the *E. coli* data for this analysis. We identified the most common patterns of occurrence among the 1,207 plasmid-bearing *E. coli* genomes, focusing on pCONJ, pMOB, pOriT and pNT (Fig 8A). The most common pattern is the presence of only conjugative plasmids in the cell. The second and fourth most frequent patterns are a pair of pCONJ-pMOB and the triplet pCONJ-pMOB-pOriT. Interestingly, the third most frequent pattern is the single presence of MOBless pNTs, in contrast to the much rarer event of having single pOriTs in the cell. This further reinforces the idea that while MOBless pNTs are non-transmissible and vertically transmitted with their host cells, pOriTs co-transfer with co-existing elements within the cell.

**Figure 8.**
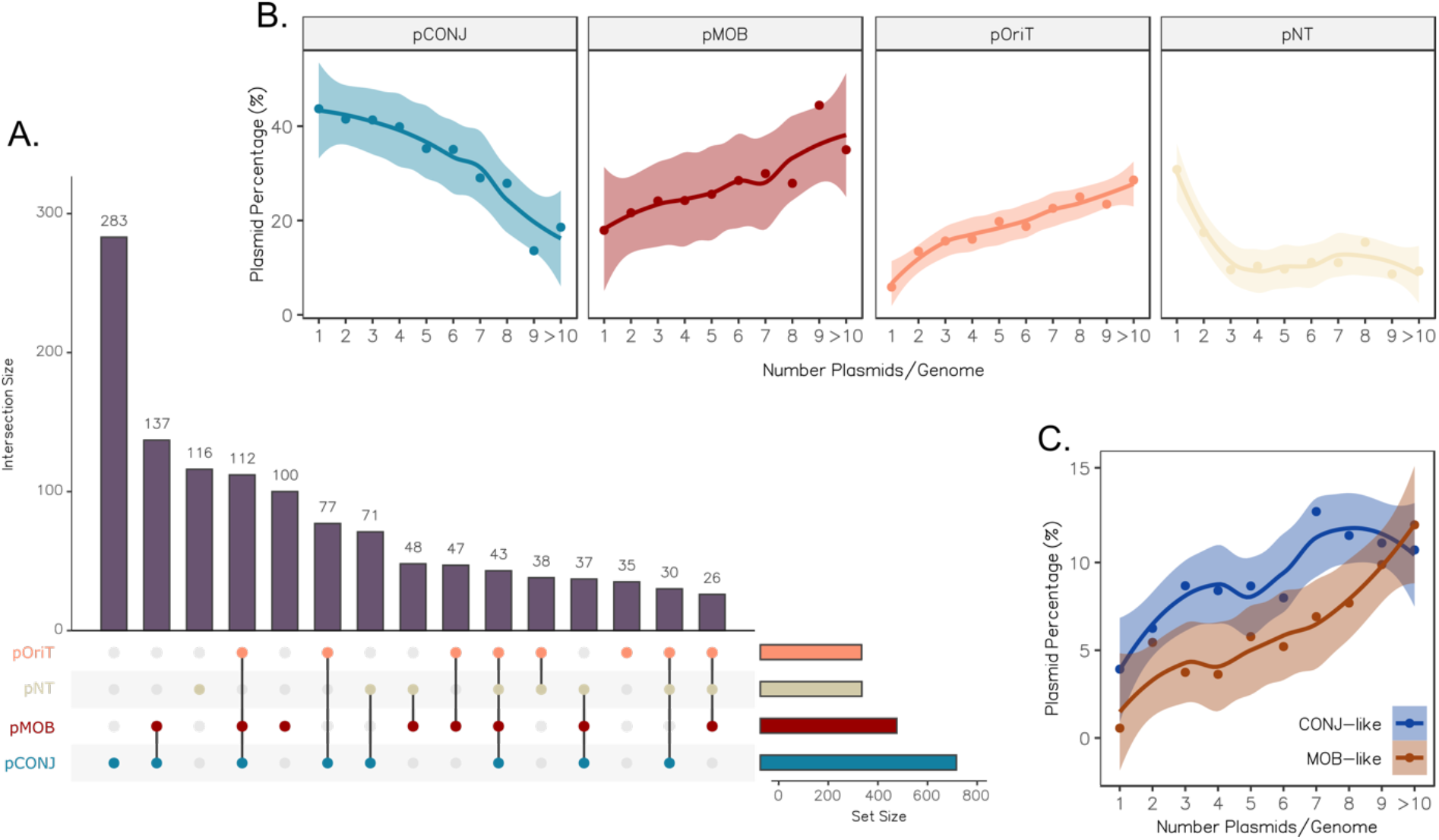
**A.** Upset plot showing the distribution of pCONJ, pMOB, pOriT and pNT co-occurrences. **B.** Proportion of plasmid types attending to the number of plasmids in their hosts’ genomes. **C.** Proportion of pCONJ-like and pMOB-like pOriTs attending to the number of plasmids in their hosts’ genomes.

If the pMOB and pOriT require a pCONJ to transfer between cells, one would expect that the frequency of each type of plasmids would vary with the number of plasmids per genome. Notably, genomes with few plasmids would tend to have more pCONJ and those with many plasmids would have progressively a larger fraction of other types of plasmids. Indeed, the frequency of pCONJ in *E. coli* is highest in genomes with a single plasmid and constantly decreases with the increase in the number of plasmids (Fig 8B). As expected, pMOB and pOriT show the inverse trend. These plasmids are rarely found alone in the genome and become increasingly frequent when cells contain more and more plasmids. The frequency of these plasmids is very high (75%) in genomes with more than 10 different plasmids. Hence, the relative frequency of each type of plasmid varies with the number of plasmids in the cell.

We showed above that some pOriTs may only require a pCONJ (since they have a pCONJ-like *oriT*), whereas others may require a pCONJ and a pMOB to transfer (pMOB-like *oriT*). The latter might be found preferentially in genomes with more plasmids, since they require a combination of two compatible plasmids to transfer. Indeed, while pCONJ-like pOriTs reach a frequency plateau in genomes with ≥7 plasmids, pMOB-like pOriTs increase steeply in frequency up to 10 plasmids/genome (Fig 8C). All these findings suggest that the functional dependencies of certain plasmids relative to others do shape the co-occurrence of plasmids in populations.

## Discussion

To understand how so many plasmids could lack relaxases and still be present across distant strains, we searched for homologs of experimentally verified *oriT*, the only genetic element a plasmid needs *in-cis* for conjugation. The search of homologs of *oriTs* could result in misidentifications, but our observations suggest that most of the *oriTs* that we identified are correct. (1) While most plasmids have an *oriT*, most chromosomes lack them, in spite of their much longer sequences. (2) At least one *oriT* has been identified in most plasmids that were expected to have it (pMOB or pCONJ). (3) There are no cross matches between *E. coli* and *S. aureus oriTs*. (4) There are almost no cross matches between pCONJs and pMOBs, allowing to identify pCONJ-like and pMOB-like *oriTs*. (5) Most plasmids have one single *oriT*, and the others often have multiple relaxases, seem to be plasmid co-integrates, or have been already described^40^. (6) Almost all *oriT*s identified are located in non-coding regions. (7) There is a strict association between the *oriTs* and their associated relaxase family. (8) The *oriTs* were not found where they were not expected, *e.g*. in phage-plasmids that rely on alternative mechanisms rather than conjugation^38^, or in pO157-like plasmids, which are known to be non-conjugative^39^. Finally, previous work in *S. aureus* validated the identification of *oriT*s in plasmids^25^. These results suggest that we identified most *oriTs* (#1, #2, #5, #6), that false positives are probably rare (#1, #3, #4, #6, #8), and that associations between *oriT* and relaxases are reliable (#4, #5, #7). Hence, our *oriT* screening seems accurate. Yet, it’s likely that some *oriTs* remain to be identified, since some pCONJ and pMOB lack known *oriT*s (Fig 2E, Fig S12). Further work will be needed to identify these novel *oriT*s across bacterial species. That will require extensive computational analysis and experimental validation of the *oriT*s representatives.

The observation that pOriTs usually have *oriT*s from either pCONJ or pMOB, suggests that these elements have evolved to either hijack the relaxase of a conjugative or a mobilizable plasmid. The latter require a pCONJ themselves resulting in a complex succession of ecological dependencies (see below). These two types of pOriT could have arisen by gene deletion of pCONJ and pMOB, in which case the pOriT would have lost the genes encoding the relaxase (and the MPF in pCONJ) while keeping the ancestral *oriT*. This is consistent with the emergence of novel pOriTs in closely related plasmids within PTUs. More complex scenarios are also possible, *e.g*. the translocation of an *oriT* to a plasmid lacking one. The hypothesis of frequent pOriT genesis by gene deletion from pMOB or pCONJ is further supported by the analysis of the distribution of pOriT size which has two modes, each slightly smaller than the modes of pMOB and pCONJ (Fig 7B, Fig S11). We have proposed that a fraction of pMOB derived recently from pCONJ^34^. Our present results further suggest that a part of pOriT originated from either pCONJ or pMOB.

Why would plasmids evolve towards less autonomous mobilization, *i. e*. to depend on other plasmids for mobility? The *oriT* is a small non-coding sequence that may have little impact on bacterial fitness. In contrast, MPF systems and relaxases are costly and may hamper the successful vertical transmission of the plasmid^41,42^. This is why the genetic components of conjugative plasmids are usually repressed^43^ and occasionally lost^44^. Hence, the loss of protein-coding genes for conjugation may decrease horizontal transfer but increase the success of vertical transmission. In contrast, the loss of *oriT*s precludes horizontal transmission by conjugation without providing significant advantages for vertical transmission. Hence, the conditions that favor loss of conjugation-related protein coding genes may not favor the loss of *oriT*.

The decrease in horizontal transmission associated with the loss of protein-coding genes for conjugation resulting in pOriT depends on the frequency with which the latter co-occurs with a compatible pCONJ (and eventually also a pMOB). We observed that the frequency of pOriT with pCONJ-like and pMOB-like *oriTs* was in direct proportion of the frequency of the “helper” plasmids. The dependence of pOriT on the presence of other plasmids in the cell might suggest that pOriTs should evolve to have a pCONJ-like *oriT* and dispense the requirement for a pMOB. Notwithstanding, pMOBs are frequent and can often be mobilized by many different pCONJ^32,33^. We speculate that pOriT with pMOB-like *oriT*s have an advantage in certain cases over those with pCONJ-like *oriTs* in that pMOB may hijack many different pCONJ. In genomes with many plasmids the right combinations pMOB/pCONJ might not be rare and allow the transfer of the pOriT. Furthermore, if the mobilization of a pOriT and/or pMOB entails the co-transfer of the helper pCONJ as it has been suggested^45^, the pOriT will find in this novel host cell all the plasmids that are required for its subsequent mobility.

Independently of the reasons leading to the high frequency of the different pOriTs, their requirements for conjugation seem to shape plasmid distribution in cells. Large and small plasmids were previously found to co-occur more often than expected in bacteria^46^. Since large plasmids are often pCONJ and smaller ones are typically pMOB or pOriT, this fits our observations of co-occurrence of the different types of plasmids. Interestingly, pMOBs and pOriTs were particularly abundant in genomes bearing many plasmids, where the chances to find helper pCONJ are high. In contrast, pCONJ, which conjugate autonomously, are the most common plasmids in cells having one or a few elements. The simplest mechanism to explain these results is that these plasmids often arrive at the cell together, *i. e*. using the same mating event. But additional interactions may also contribute to further stabilize the presence of these plasmids in cells. For example, the cost of carrying small plasmids was smaller in a *Pseudomonas* strain already carrying a large plasmid^46^.

Our results suggest that the majority of plasmids are able to conjugate autonomously or by recruitment of functions from other plasmids. Considering classical and RCR-mediated conjugation, around 90% of *S. aureus* plasmids have the genetic elements needed to be horizontally transferred via conjugation. Notwithstanding, alternative mechanisms of plasmid mobility have been recently described. Among *E. coli* plasmids, there are 7% of phage-plasmids that can transfer within their own viral particles. In *S. aureus*, phage-plasmids are rare, but plasmids can be transduced by phages and their satellites^47^. Phages and satellites can transduce pieces of DNA of approximately the size of their own genomes. The size of the genomes of temperate phages matches the largest mode of the sizes of pMOBless and the size of the satellite genomes matches the smallest mode of these plasmids. It was proposed that plasmids were selected to have sizes compatible with transduction by phages and satellites, which explains the bi-modal distribution of plasmid sizes (Fig 7B)^47^. If correct, transduction by phages and their satellites would explain the enigmatic bi-modality of plasmid sizes, while gene deletions causing the transitions between pCONJ or pMOB to pOriT would explain why the latter tend to follow the size distribution of the former.

In summary, 9 out of 10 plasmids bear identifiable genetic elements that may mediate their horizontal transfer, most of them by conjugation. There are only ~10% plasmids lacking known genetic elements associated with horizontal transfer. Such plasmids may still occasionally be transferred through alternative mechanisms leaving little trace in the plasmid sequence, such as transformation or transduction. With this work, we provide strong evidence suggesting that there is no conundrum regarding the plasmid mobility, and provide new insights into alternative mechanisms of plasmid transfer.

## Methods

### Genome data

We retrieved from all the complete genomes available in the NCBI non-redundant RefSeq database in March 2021 (22,255 genomes, 21,520 plasmids) those of *Escherichia coli* and *Staphylococcus aureus* species. These resulted in a set of 1,585 genomes of *Escherichia coli* and 582 genomes of *Staphylococcus aureus*, including 3,409 and 462 plasmids, respectively. The accession numbers and further information on the plasmids is available in the Supplementary Table 1. The information on the chromosomes and the relevant data is available on the Supplementary Table 2.

### Collection of the *oriT* database and its identification in the complete genomes

We built a collection of experimentally validated origins of transfer. First, we retrieved the 52 *oriTs* with a status *‘experimental’* from the already published *oriT* database by Li and collaborators^31^. We expanded this collection by consulting the literature, using as a query *“oriT”* in the PubMed database (available in September 2021). Among the 708 entries, we screened for experimentally validated *oriT*s not included in the aforementioned database. This resulted in the retrieval of 47 additional *oriT*s. However, 1 *oriT* from the published database and 7 *oriT*s from the literature were discarded from the collection as only the *nic*-site sequence was available. This resulted in a final dataset of 91 origins of transfer. Information on this collection is available in Supplementary Table 3.

We used BLAST, version 2.9.0+, to identify *oriT*s^48^. The complete genomes of *E. coli* and *S. aureus* were indexed with makeblastdb. Then, we used blastn to search for occurrences of each of the 91 *oriTs* (query) against the database of complete genomes. Due to the short length of the origins of transfer, blastn was used with the option *-task blastn-short* and an E-value threshold of 0.01 following the developer’s instructions. In cases in which two different *oriTs* were identified in the same region of a plasmid (overlapping), the *oriT* hit with the best E-value was retrieved.

We identified during this screening an exceptional case of a ~50 kb plasmid with 23 identical *oriTs*.This plasmid (NZ_CP019265.1) was discarded from further analysis as we considered it to be a sequencing artifact.

### Characterization of conjugative systems and relaxases and plasmid classification on the mobility

We used the module CONJscan of MacSyFinder, version 2.0^49^ to identify all the complete MPF systems. The individual hidden Markov model (HMM) hits that were not associated with MPFs deemed complete were used to identify incomplete MPF systems.

Relaxases were identified using HMMER version 3.3.2^50^, and the HMM profiles employed by the software MOBscan^51^. We used the tool hmmsearch (default options) to screen for relaxases in all the proteins annotated in the dataset and kept the 2,195 significant hits with >50% coverage on the profile. A careful analysis of the results revealed that this version of the RefSeq annotations sometimes missed genes encoding relaxases, especially when these genes overlapped others (Fig S13). To correct for this artifact, we introduced a preliminary step of re-annotation to ensure a coherent annotation of the genes throughout all the genomes, which was then used to identify the MPF and the relaxases. For the annotation, we used the software Prodigal, version 2.6.3^52^, with the recommended mode for plasmids and viruses to identify all open reading frames. Hits were then identified as mentioned above. When two different profiles matched the same protein, we kept the one with the lowest E-value.

Following the previous characterization, plasmids were classified in different mobility categories depending on their composition in terms of *oriT*, relaxase, and MPF genes. Plasmids encoding a putatively complete MPF system (including a relaxase) were considered to be conjugative (pCONJ). Plasmids encoding relaxases and lacking a complete MPF system were classified as mobilizable (pMOB). The remaining plasmids were classified as pMOBless, and were split into different categories: pOriTs when they had an *oriT*, phage-plasmids (P-Ps) when they were phage-related elements (see below) or presumably non-transmissible (pNTs) otherwise. In addition, some plasmids were classified as decayed conjugative plasmids (pdCONJ). These plasmids encode two or more MPF genes, but not enough to form a complete MPF system. Therefore, pdCONJ show a close evolutionary relationship with conjugative plasmids^34^, but are considered pMOB, pOriT or pNT in terms of mobility (Fig S14). Similarly, the loci encoding presumably complete MPF systems in chromosomes were classed as ICE (Integrative and Conjugative Element), even if often we ignore the precise limits of the element. Chromosomal genes encoding relaxases that were distant from genes encoding MPFs (> 60 genes) were classed as IME (Integrative and Mobilizable Element).

### Identification of Rolling Circle Replication Proteins

For the identification of Rolling Circle Replication (RC-Rep) proteins involved in plasmid conjugation, we first retrieved the RC-Rep of the *Staphylococcus aureus* plasmid pC194 (NC_002013.1), a pNT plasmid known to be mobilized through *in trans* conjugation^36^. We used its Pfam profile^53^, Rep_1 (PF01446), to look for related RC-Rep proteins in all the plasmids of *E. coli* and *S. aureus* using the HMMER tool hmmsearch (default options, E-value < 0.001), version 3.3.2^50^.

### Identification of phage-plasmids

For the identification of phage-plasmids (P-Ps), we retrieved the *E. coli* and *S. aureus* P-Ps recently unveiled^54^. The database used in the cited work corresponds to the same RefSeq database (retrieved on March 2021). This way, we were able to identify 222 P-Ps among the 3,409 *E. coli* plasmids and 1 P-P among the 482 *S. aureus* plasmids.

### Analysis of the pangenome of *E. coli* and *S. aureus* plasmids

The pangenome of the plasmid-encoded genes of *E. coli* and *S. aureus* was identified using the module pangenome of the software PanACoTa, version 1.3.1^55^. Briefly, gene families were built with MMseqs2, version 13.45111, with an identity threshold of 80%. This is the typical threshold for the determination of the *E. coli* pangenome^30^. This way, the 227,428 plasmid-encoded proteins in *E. coli* were grouped into 11,530 gene families. In *S. aureus*, the 7,902 proteins were grouped into 1,010 gene families. Some plasmids were not used in the analysis because their annotations lacked protein coding genes: 32 of the 3,409 plasmids in *E. coli* (0.94%) and 20 of the 482 in *S. aureus* (4,15%). Rarefaction curves were performed with the R package vegan, version 2.5-6^57^. The later package was additionally employed to infer the plasmid pangenome of *S. aureus* until matching the same sample size as *E. coli* following an Arrhenius model. Additionally, the Gleason model and Gitay model were used to extrapolate the rarefaction curves of the pangenome for *S. aureus* (Fig S15). Rarefaction curves were plotted with sample sizes increasing by a step of 100 plasmids.

### Determination of sequence similarity between plasmids

We assessed sequence similarity for all pairs of the 3,869 plasmids using two different approaches.

To analyze very closely related plasmids, we classified them based on their average nucleotide identity (ANI) into the existing catalogue of Plasmid Taxonomic Units (PTUs)^27^. The clustering was performed using COPLA^58^, version 1.0 (default parameters).

To analyze more distantly related plasmids, we assessed the gene relatedness within and between PTUs, using the weighted Gene Repertoire Relatedness (wGRR)^59^. For this, we searched for sequence similarity between all the proteins identified in the plasmids using MMseqs2 (version 9-d36de)^56^, retrieving the hits with E-value < 10^−4^ and coverage > 50%. Best bi-directional hits (BBH) between pairs of plasmids were used to calculate the wGRR as previously described^59^:

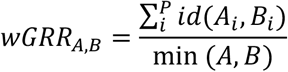

where *A*_i_ and *B_i_* are the *i*th BBH pair of *P* total pairs; *id*(*A_i_, B_i_*) is the identity between the BBH pair; and *min*(*A, B*) is the number of genes encoded in the smallest plasmid of the pair. This way, the wGRR value varies between 0 (no BBH between the plasmids) and 1 (all genes of the smallest plasmid have an identical homolog in the larger one). The wGRR values were used to identify related plasmids between and within PTUs, setting the threshold in wGRR > 0.75 as previously described^34^. With this purpose, only plasmid pairs with wGRR > 0.75 were retrieved for visualizations, *i.e*. at least the 75% of genes encoded in the smallest plasmid are shared between the pair.

### Clustering of the *oriTs*

We clustered the *oriTs* in families, by searching for sequence similarity between all pairs of *oriTs* in the reference dataset using blastn^48^ (Fig S16). BLAST was used with the option *-task blastn-short* and an E-value threshold of 0.01. Only matches with >80% identity and >70% coverage of the smallest *oriT* were kept for the clustering analysis. The clustering was performed with the hierarchical method available in the R package pheatmap, version 1.0.12 (default options)^60^. The clusters were named after well-known *oriT*s contained in the cluster: F-like, R6K-like, R64-like, ColE1-like, RP4-like and R46-like. The association of each *oriT* to their *oriT* family is available in the Supplementary Table 3 and Supplementary Figure 16.

### Determination of antimicrobial resistance genes

For the identification of antimicrobial resistance genes encoded in the plasmid dataset, we used AMRFinderPlus^61^, version 3.10, with the default options. This tool combines BLASTP and HMMER to identify the 6,189 resistance determinants available in the NCBI Pathogen Detection Reference Gene Catalog (April 2022). The latter is the result of the curated merging of various widespread-used databases, including CARD^62^, and ResFinder^63^ databases, among others^61^.

### Statistical analysis

Except where explicitly stated, all statistical analyses were done with R, version 3.5.2. Additionally, all visualizations were performed with the R package ggplot2^64^, version 3.3.5, occasionally supported by the R packages ggsignif^65^, version 0.6.0 and ggridges^66^, version 0.5.3. For the construction and visualization of the networks, we used the R package igraph^67^, version 1.2.4.1 and the software Gephi 0.9.2^68^, respectively.

## Acknowledgements

We would like to thank Eugen Pfeifer for providing the wGRR and PTUs data, Fernando de la Cruz and Maria Pilar Garcillán Barcia for discussion along the years on plasmid mobility. Microbial Evolutionary Genomics Unit for scientific discussions. INCEPTION project [PIA/ANR-16-CONV-0005]. Fédération pour la Recherche Médicale [Equipe FRM/EQU201903007835]. Labex IBEID [ANR-10-LABX-62-IBEID]. HORIZON-MSCA-2021-PF-01-01 EvoPlas-101062386 to Manuel Ares-Arroyo.

## Competing interests

The authors declare no competing interests.

## Notes

### Competing Interest Statement

The authors have declared no competing interest.

## References

1. Treangen, T. J. & Rocha, E. P. C. Horizontal Transfer, Not Duplication, Drives the Expansion of Protein Families in Prokaryotes. PLoS Genet 7, e1001284 (2011).

2. Frost, L. S., Leplae, R., Summers, A. O. & Toussaint, A. Mobile genetic elements: the agents of open source evolution. Nat Rev Microbiol 3, 722–732 (2005).

3. Wein, T. & Dagan, T. Plasmid evolution. Curr Biol 30, R1158–R1163 (2020).

4. Zhang, X. et al. Improvement in the efficiency of natural transformation of *Haemophilus parasuis* by shuttle-plasmid methylation. Plasmid 98, 8–14 (2018).

5. Erdmann, S., Tschitschko, B., Zhong, L., Raftery, M. J. & Cavicchioli, R. A plasmid from an Antarctic haloarchaeon uses specialized membrane vesicles to disseminate and infect plasmid-free cells. Nat Microbiol 2, 1446–1455 (2017).

6. Canosi, U., Lüder, G. & Trautner, T. A. SPP1-mediated plasmid transduction. J Virol 44, 431–436 (1982).

7. Pfeifer, E., Moura de Sousa, J. A., Touchon, M. & Rocha, E. P. C. Bacteria have numerous distinctive groups of phage–plasmids with conserved phage and variable plasmid gene repertoires. Nucleic Acids Res 49, 2655–2673 (2021).

8. Smillie, C., Garcillán-Barcia, M. P., Francia, M. V., Rocha, E. P. C. & de la Cruz, F. Mobility of Plasmids. Microbiol Mol Biol Rev 74, 434–452 (2010).

9. De La Cruz, F., Frost, L. S., Meyer, R. J. & Zechner, E. L. Conjugative DNA metabolism in Gram-negative bacteria. FEMS Microbiol Rev 34, 18–40 (2010).

10. Guglielmini, J., de la Cruz, F. & Rocha, E. P. C. Evolution of Conjugation and Type IV Secretion Systems. Mol Biol Evol 30, 315–331 (2013).

11. Johnson, C. M. & Grossman, A. D. Integrative and Conjugative Elements (ICEs): What They Do and How They Work. Annu. Rev. Genet. 49, 577–601 (2015).

12. Cury, J., Oliveira, P. H., de la Cruz, F. & Rocha, E. P. C. Host Range and Genetic Plasticity Explain the Coexistence of Integrative and Extrachromosomal Mobile Genetic Elements. Mol Biol Evol 35, 2230–2239 (2018).

13. Branger, C. et al. Specialization of small non-conjugative plasmids in *Escherichia coli* according to their family types. Microbial Genom 5, e000281 (2019).

14. Gu, D. et al. A fatal outbreak of ST11 carbapenem-resistant hypervirulent *Klebsiella pneumoniae* in a Chinese hospital: a molecular epidemiological study. Lancet Infect Dis 18, 37–46 (2018).

15. Maeda, S. et al. Horizontal transfer of nonconjugative plasmids in a colony biofilm of *Escherichia coli*. FEMS Microbiol Lett 255, 115–120 (2006).

16. Lambert, C. M., Hyde, H. & Strike, P. Conjugal mobility of the multicopy plasmids NTP1 and NTP16. Plasmid 18, 99–110 (1987).

17. Chang, A. C. & Cohen, S. N. Construction and characterization of amplifiable multicopy DNA cloning vehicles derived from the P15A cryptic miniplasmid. J Bacteriol 134, 1141–1156 (1978).

18. Lee, C. A., Thomas, J. & Grossman, A. D. The *Bacillus subtilis* Conjugative Transposon ICE *Bs1* Mobilizes Plasmids Lacking Dedicated Mobilization Functions. J Bacteriol 194, 3165–3172 (2012).

19. Xie, M. et al. Conjugation of Virulence Plasmid in Clinical *Klebsiella pneumoniae* Strains through Formation of a Fusion Plasmid. Adv Biosyst 4, e1900239 (2020).

20. Daccord, A., Ceccarelli, D. & Burrus, V. Integrating conjugative elements of the SXT/R391 family trigger the excision and drive the mobilization of a new class of *Vibrio* genomic islands: ICE-mediated GI mobilization. Mol Microbiol 78, 576–588 (2010).

21. Brockhurst, M. A. & Harrison, E. Ecological and evolutionary solutions to the plasmid paradox. Trends Microbiol 30, 534–543 (2022).

22. Hall, J. P. J., Wood, A. J., Harrison, E. & Brockhurst, M. A. Source–sink plasmid transfer dynamics maintain gene mobility in soil bacterial communities. Proc. Natl. Acad. Sci. U.S.A. 113, 8260–8265 (2016).

23. O’Brien, F. G. et al. Origin-of-transfer sequences facilitate mobilisation of non-conjugative antimicrobial-resistance plasmids in *Staphylococcus aureus*. Nucleic Acids Res 43, 7971–7983 (2015).

24. Pollet, R. M. et al. Processing of Nonconjugative Resistance Plasmids by Conjugation Nicking Enzyme of Staphylococci. J Bacteriol 198, 888–897 (2016).

25. Ramsay, J. P. et al. An updated view of plasmid conjugation and mobilization in *Staphylococcus*. Mob Genet Elements 6, e1208317 (2016).

26. Ramsay, J. P. & Firth, N. Diverse mobilization strategies facilitate transfer of non-conjugative mobile genetic elements. Curr Opin Microbiol 38, 1–9 (2017).

27. Redondo-Salvo, S. et al. Pathways for horizontal gene transfer in bacteria revealed by a global map of their plasmids. Nat Commun 11, 3602 (2020).

28. Murray, C. J. et al. Global burden of bacterial antimicrobial resistance in 2019: a systematic analysis. Lancet 12, 629–655 (2022).

29. Partridge, S. R., Kwong, S. M., Firth, N. & Jensen, S. O. Mobile Genetic Elements Associated with Antimicrobial Resistance. Clin Microbiol Rev 31, e00088–17 (2018).

30. Touchon, M. et al. Phylogenetic background and habitat drive the genetic diversification of *Escherichia coli*. PLoS Genet 16, e1008866 (2020).

31. Li, X. et al. oriTfinder: a web-based tool for the identification of origin of transfers in DNA sequences of bacterial mobile genetic elements. Nucleic Acids Res 46, W229–W234 (2018).

32. Cabezón, E., Lanka, E. & de la Cruz, F. Requirements for mobilization of plasmids RSF1010 and ColE1 by the IncW plasmid R388: trwB and RP4 traG are interchangeable. J Bacteriol 176, 4455–4458 (1994).

33. Sastre, J. I., Cabezón, E. & de la Cruz, F. The carboxyl terminus of protein TraD adds specificity and efficiency to F-plasmid conjugative transfer. J Bacteriol 180, 6039–6042 (1998).

34. Coluzzi, C., Garcillán-Barcia, M. P., de la Cruz, F. & Rocha, E. P. C. Evolution of Plasmid Mobility: Origin and Fate of Conjugative and Nonconjugative Plasmids. Mol Biol Evol 39, msac115 (2022).

35. Moran, R. A. & Hall, R. M. Analysis of pCERC7, a small antibiotic resistance plasmid from a commensal ST131 *Escherichia coli*, defines a diverse group of plasmids that include various segments adjacent to a multimer resolution site and encode the same NikA relaxase accessory protein enabling mobilisation. Plasmid 89, 42–48 (2017).

36. Lee, C. A., Thomas, J. & Grossman, A. D. The *Bacillus subtilis* conjugative transposon ICEBs1 mobilizes plasmids lacking dedicated mobilization functions. J Bacteriol 194, 3165–3172 (2012).

37. Garcillán-Barcia, M. P., Pluta, R., Lorenzo-Díaz, F., Bravo, A. & Espinosa, M. The Facts and Family Secrets of Plasmids That Replicate via the Rolling-Circle Mechanism. Microbiol Mol Biol Rev 86, e0022220 (2022).

38. Łobocka, M. B. et al. Genome of bacteriophage P1. J Bacteriol 186, 7032–7068 (2004).

39. Lim, J. Y., Yoon, J. & Hovde, C. J. A brief overview of *Escherichia coli* O157:H7 and its plasmid O157. J Microbiol Biotechnol 20, 5–14 (2010).

40. Avila, P., Núñez, B. & de la Cruz, F. Plasmid R6K contains two functional *oriTs* which can assemble simultaneously in relaxosomes in vivo. J Mol Biol 261, 135–143 (1996).

41. San Millan, A. & MacLean, R. C. Fitness Costs of Plasmids: a Limit to Plasmid Transmission. Microbiol Spectr 5 (2017).

42. Turner, P. E., Cooper, V. S. & Lenski, R. E. TRADEOFF BETWEEN HORIZONTAL AND VERTICAL MODES OF TRANSMISSION IN BACTERIAL PLASMIDS. Evolution 52, 315–329 (1998).

43. Koraimann, G. & Wagner, M. A. Social behavior and decision making in bacterial conjugation. Front Cell Infect Microbiol 4, 54 (2014).

44. Hooton, S. P. T. et al. Laboratory Stock Variants of the Archetype Silver Resistance Plasmid pMG101 Demonstrate Plasmid Fusion, Loss of Transmissibility, and Transposition of Tn7/pco/sil Into the Host Chromosome. Front Microbiol 12, 723322 (2021).

45. Dionisio, F., Zilhão, R. & Gama, J. A. Interactions between plasmids and other mobile genetic elements affect their transmission and persistence. Plasmid 102, 29–36 (2019).

46. San Millan, A., Heilbron, K. & MacLean, R. C. Positive epistasis between co-infecting plasmids promotes plasmid survival in bacterial populations. ISME J 8, 601–612 (2014).

47. Humphrey, S. et al. Staphylococcal phages and pathogenicity islands drive plasmid evolution. Nat Commun 12, 5845 (2021).

48. Camacho, C. et al. BLAST+: architecture and applications. BMC Bioinformatics 10, 421 (2009).

49. Cury, J., Abby, S. S., Doppelt-Azeroual, O., Néron, B. & Rocha, E. P. C. Identifying Conjugative Plasmids and Integrative Conjugative Elements with CONJscan. Methods Mol Biol 2075, 265–283 (2020).

50. Wheeler, T. J. & Eddy, S. R. nhmmer: DNA homology search with profile HMMs. Bioinformatics 29, 2487–2489 (2013).

51. Garcillán-Barcia, M. P., Redondo-Salvo, S., Vielva, L. & de la Cruz, F. MOBscan: Automated Annotation of MOB Relaxases. Methods Mol Biol 2075, 295–308 (2020).

52. Hyatt, D. et al. Prodigal: prokaryotic gene recognition and translation initiation site identification. BMC Bioinformatics 11, 119 (2010).

53. Mistry, J. et al. Pfam: The protein families database in 2021. Nucleic Acids Res 49, D412–D419 (2021).

54. Pfeifer, E., Bonnin, R. & Rocha, E. P. C. Phage-plasmids spread antibiotic resistance genes through infection and lysogenic conversion. http://biorxiv.org/lookup/doi/10.1101/2022.06.24.497495 (2022).

55. Perrin, A. & Rocha, E. P. C. PanACoTA: a modular tool for massive microbial comparative genomics. NAR Genom Bioinform 3, lqaa106 (2021).

56. Steinegger, M. & Söding, J. MMseqs2 enables sensitive protein sequence searching for the analysis of massive data sets. Nat Biotechnol 35, 1026–1028 (2017).

57. Okansen, J. et al. vegan: Community Ecology Package. R package version 2.5-6. https://CRAN.R-project.org/package=vegan (2019).

58. Redondo-Salvo, S. et al. COPLA, a taxonomic classifier of plasmids. BMC Bioinformatics 22, 390 (2021).

59. Cury, J., Touchon, M. & Rocha, E. P. C. Integrative and conjugative elements and their hosts: composition, distribution and organization. Nucleic Acids Res 45, 8943–8956 (2017).

60. Kolde, R. pheatmap: Pretty Heatmaps. R package version 1.0.12. https://CRAN.R-project.org/package=pheatmap (2019).

61. Feldgarden, M. et al. AMRFinderPlus and the Reference Gene Catalog facilitate examination of the genomic links among antimicrobial resistance, stress response, and virulence. Sci Rep 11, 12728 (2021).

62. Alcock, B. P. et al. CARD 2020: antibiotic resistome surveillance with the comprehensive antibiotic resistance database. Nucleic Acids Res 48, D517–D525 (2020).

63. Bortolaia, V. et al. ResFinder 4.0 for predictions of phenotypes from genotypes. J Antimicrob Chemother 75, 3491–3500 (2020).

64. Wickham, H. ggplot2: Elegant Graphics for Data Analysis. (Springer-Verlag New York, 2016).

65. Ahlmann-Eltze, C. ggsignif: Significance Brackets for ‘ggplot2’. R package version 0.6.0. https://CRAN.R-project.org/package=ggsignif (2019).

66. Wilke, C. O. ggridges: Ridgeline Plots in ‘ggplot2’. R package version 0.5.3. https://CRAN.R-project.org/package=ggridges (2021).

67. Csárdi, G. & Nepusz, T. The igraph software package for complex network research. Int J Complex Syst 1695, 1–9 (2006).

68. Bastian, M., Heymann, S. & Jacomy, M. Gephi: An Open Source Software for Exploring and Manipulating Networks. ICWSM 3, 361–362 (2009).

