## Supplementary Files for "Towards solving the conundrum of plasmid mobility: networks of functional dependencies shape plasmid transfer"

Manuel Ares-Arroyo, Charles Coluzzi, and Eduardo P.C. Rocha
Institut Pasteur, Université de Paris Cité, CNRS UMR3525, Microbial Evolutionary Genomics, Paris, France

**Table of Contents:**

**Figure S1**. *E.* coli and *S. aureus*' plasmid repertoires. [Page 2]

**Figure S2**. Analysis of the origin of transfer reference dataset. [Page 3]

**Figure S3**. Identification of *oriT*s. Page 2. [Page 4]

**Figure S4**. Chromosomal *oriT*s. [Page 5]

**Figure S5**. PTUs of the plasmids. [Page 6]

**Figure S6**. Analysis of the conjugative PTU-C. [Page 7]

**Figure S7**. Analysis of the mobilizable PTU-E22. [Page 8]

**Figure S8**. Identification of RC-Rep proteins in *S. aureus*. [Page 9]

**Figure S9**. Identification of phage-plasmids (P-Ps). [Page 10]

**Figure S10**. Analysis of putatively non-transmissible plasmids (pNTs). [Page 11]

**Figure S11**. Plasmid size according to the mobility. [Page 12]

**Figure S12**. PTUs with non-identified *oriTs*. [Page 13]

**Figure S13**. Relaxase identification. [Page 14]

**Figure S14**. Decayed conjugative plasmids. [Page 15]

**Figure S16**. Pangenome analysis of *E. coli* and *S. aureus* plasmids. [Page 16]

**Figure S16**. *oriT* families among the origins of transfer identified in the plasmid collection. [Page 17]

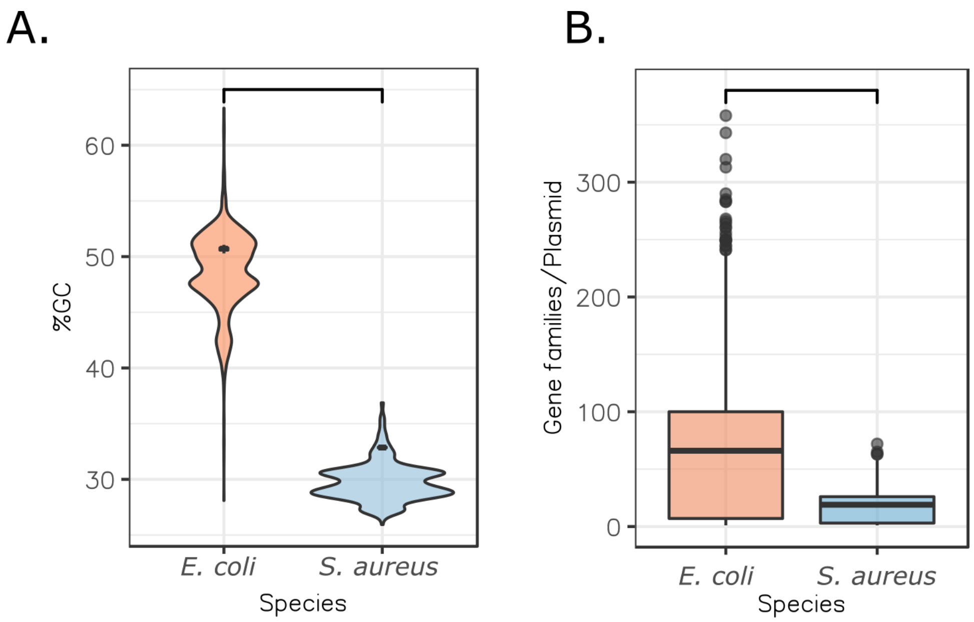


**Supplementary Figure 1. *E. coli* and *S. aureus*' plasmid repertoires. A.** Plasmid %GC. The black dots inside the violin plots represent the average GC content of the chromosomes for the set of genomes with these plasmids. The horizontal bar over the plot denotes a statistically significant difference (Student's t-test, $p < 0.0001$). **B.** Number of gene families per plasmid. The horizontal bar over the plot denotes statistically significant difference (Student's t-test, $p < 0.0001$).

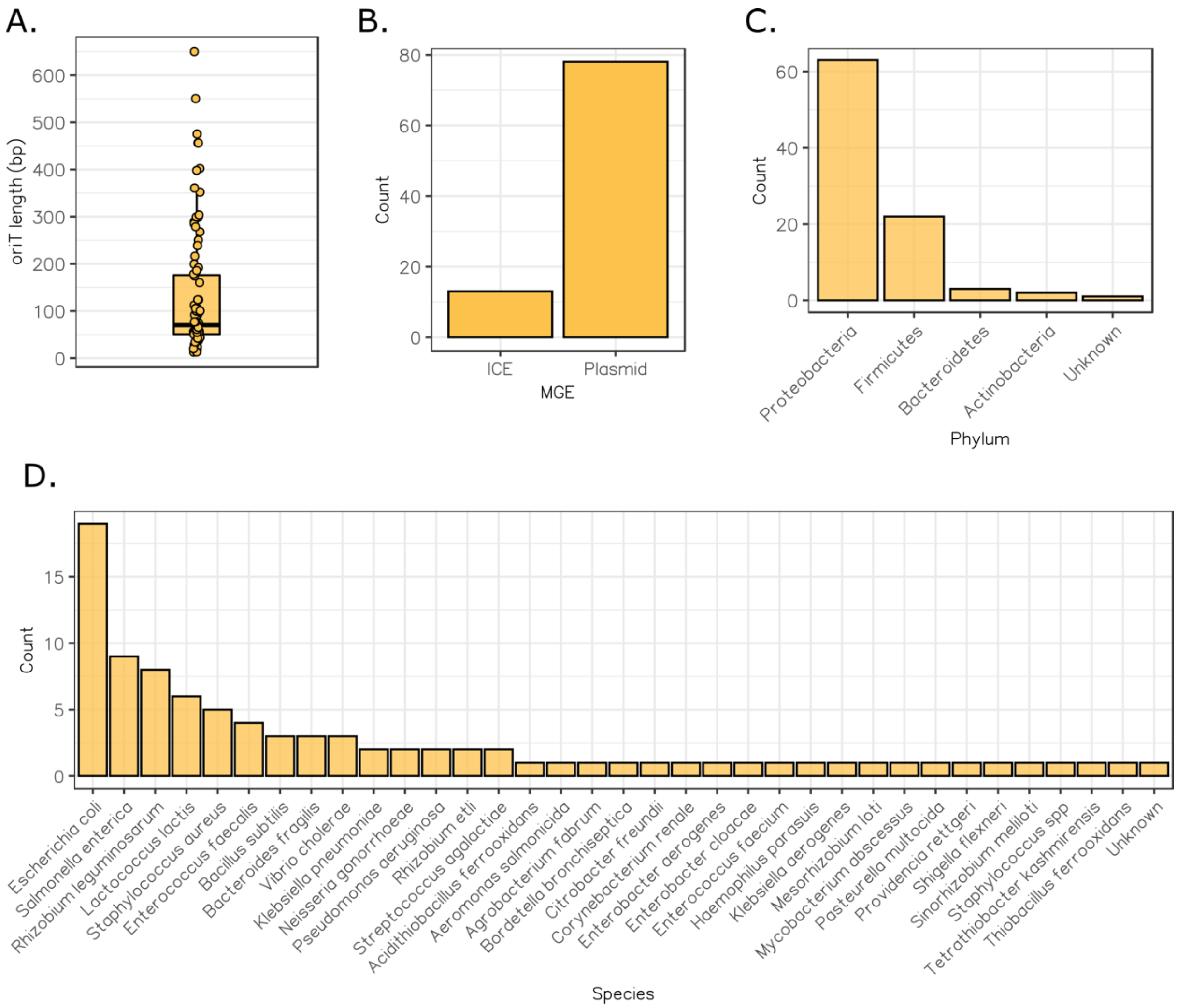


**Supplementary Figure 2. Analysis of the origin of transfer reference dataset. A.** Size of the *oriTs* included in the database. **B.** Mobile genetic element (MGE) in which the *oriTs* were first identified. ICE: Integrative and Conjugative Element. **C.** Phylum in which the *oriTs* were first described. **D.** Species in which the *oriTs* were first described.

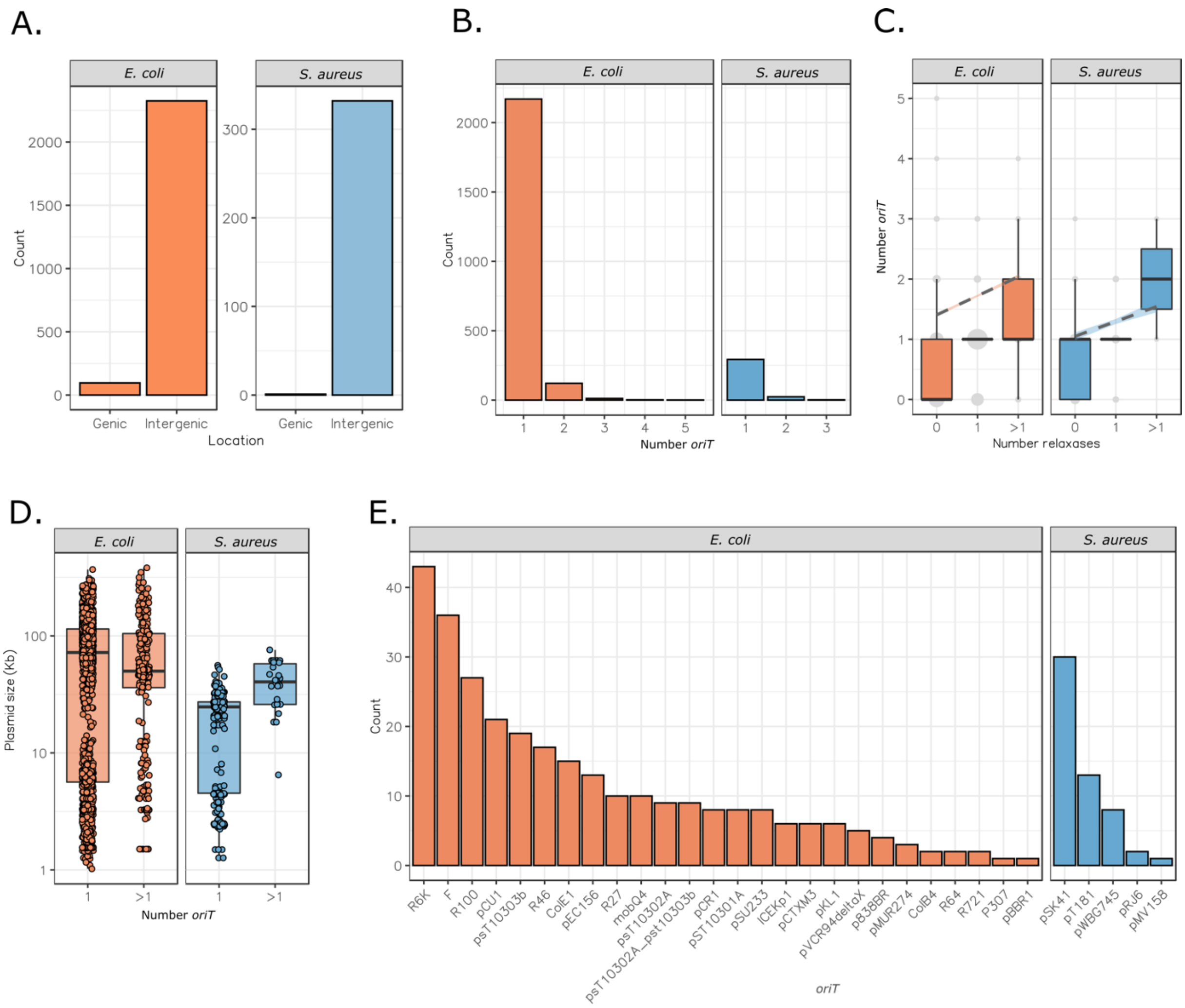


**Supplementary Figure 3. Identification of *oriT*s**. **A.** Genetic location of the *oriT*s identified, either genic or intergenic. **B.** Number of *oriTs* identified in plasmids. **B.** Number of *oriTs* according to the number of relaxases identified in the plasmid. **C.** Size of the plasmids encoding a unique *oriT* and multiple *oriTs*, respectively. **D.** *oriTs* identified in plasmids encoding multiple *oriTs*.

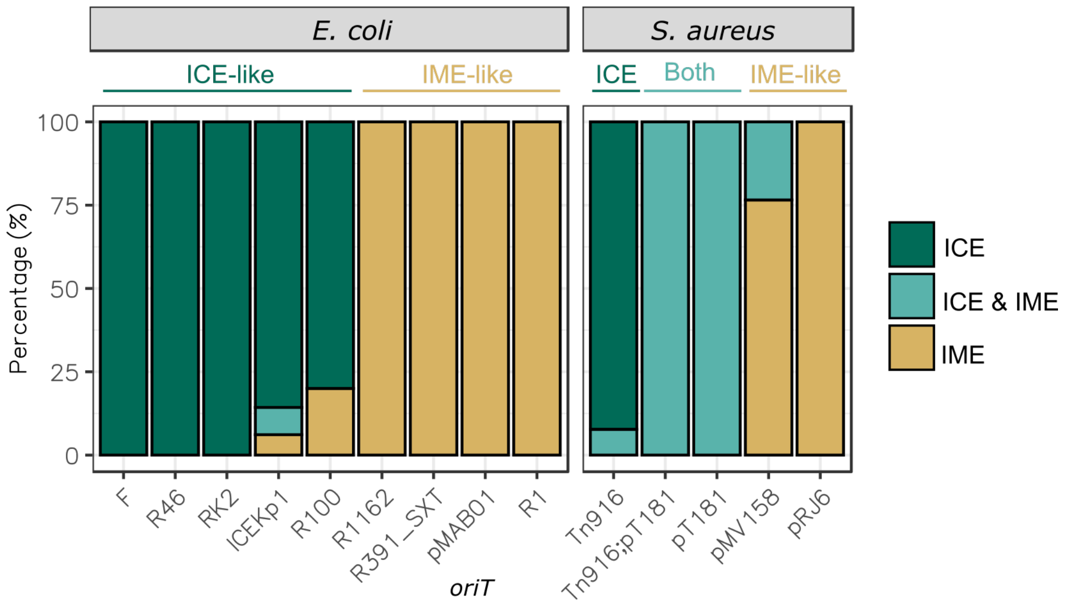


**Supplementary Figure 4. Chromosomal *oriTs*.** Proportion of integrative elements containing a given *oriT*. Analysis done for *oriT*s found in more than 5 chromosomes. ICE-like: *oriT* identified mostly in ICEs; IME-like: *oriT* identified mostly in IMEs; Unsp.: *oriT* identified in genomes with both ICEs and IMEs.

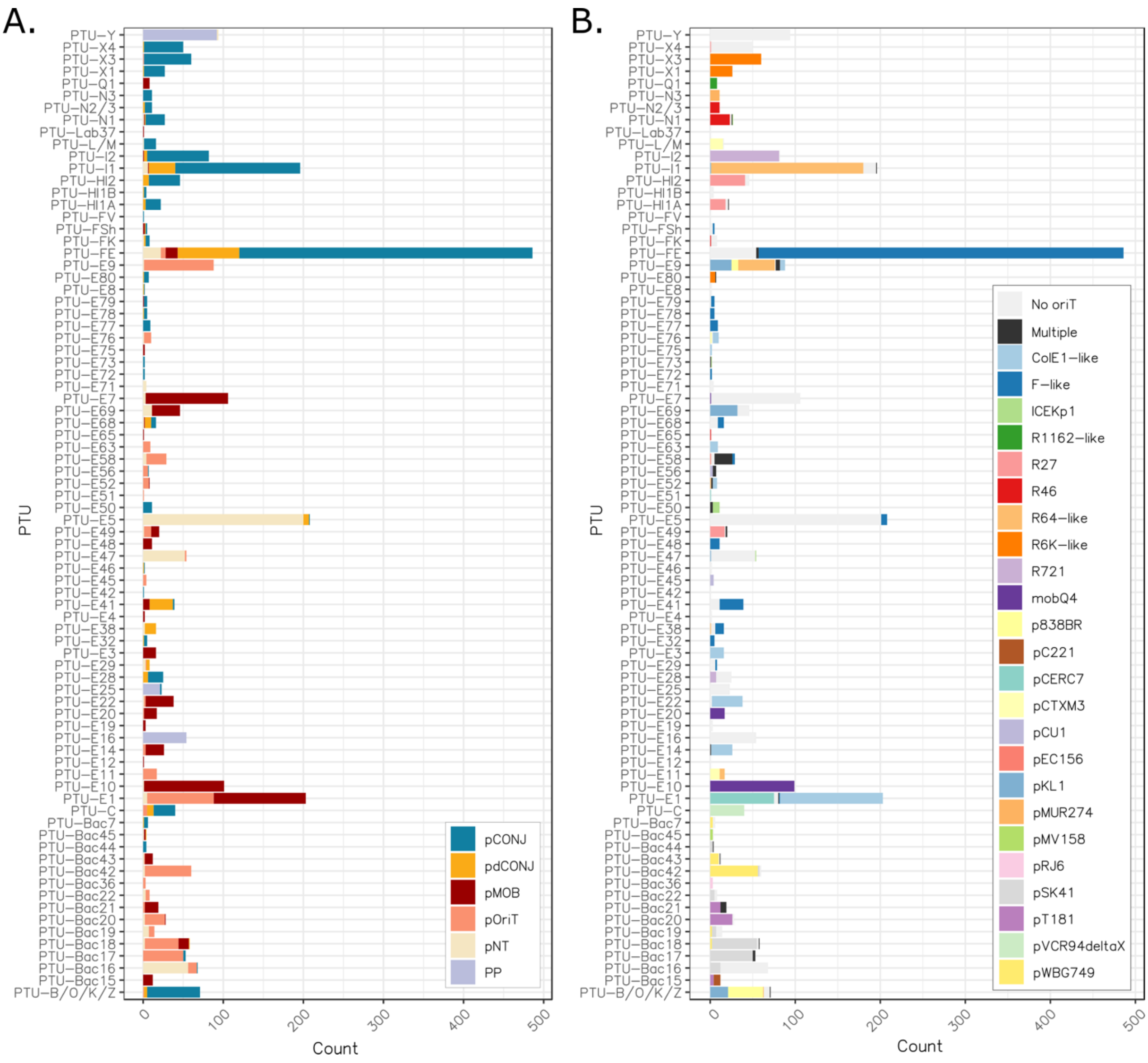


**Supplementary Figure 5. PTUs of the plasmids. A.** Colors denote the mobility of plasmids assigned to the PTUs. **B.** Colors denote the *oriTs* associated to the plasmids.

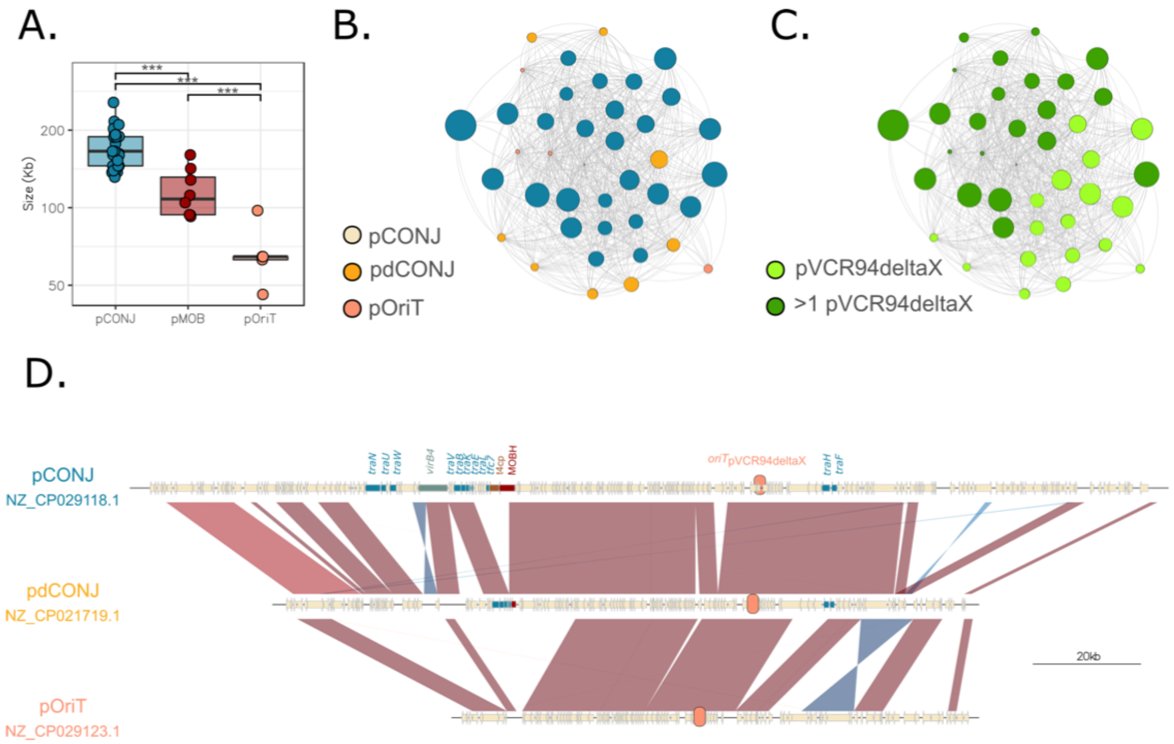


**Supplementary Figure 6. Analysis of the conjugative PTU-C. A.** Plasmid size of the PTU C according to their mobility. The horizontal bars over the plot denote statistically significant differences (pairwise t-tests, $p<0.001$). **B.** and **C.** Graph showing the PTU C. Nodes represent the plasmids and edges connect plasmid pairs with wGRR>0.75. The colors of the nodes represent the plasmid mobility type (**B**) and the *oriT* (**C**). **D.** An example of the multiple alignment of a pCONJ, pdCONJ, and pOriT from the PTU C. Genes are indicated as arrows with the relaxase in red, the coupling protein in brown, *virB4* in green, other conjugative genes in blue and the *oriT* as an orange circle.

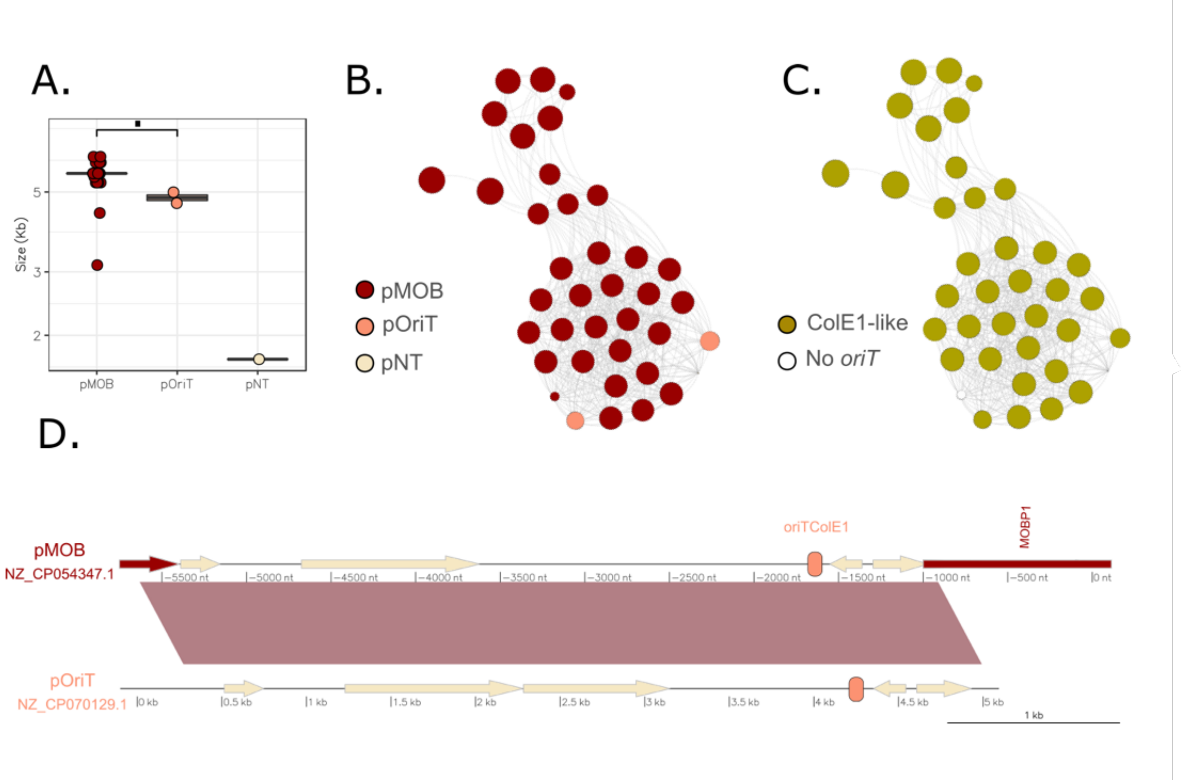


**Supplementary Figure 7. Analysis of the mobilizable PTU-E22. A.** Size of plasmids within the PTU E22 classed according to their mobility. The horizontal bar over the plot denotes the statistically significant difference (pairwise t-tests, $p<0.05$). The pNT plasmid, despite considerably smaller (<2kb) than the pMOB and pOriT, could not be tested since it has only one representative. **B.** and **C.** Graph showing the PTU E22. Nodes represent the plasmids and edges connect plasmid pairs with wGRR>0.75. The colors of the nodes represent the plasmid mobility (**B.**) and the *oriT* (**C.**). **D.** Plasmid alignments of a pMOB and pOriT from the PTU E22. The relaxase is represented as a red arrow, and the *oriT* as an orange circle.

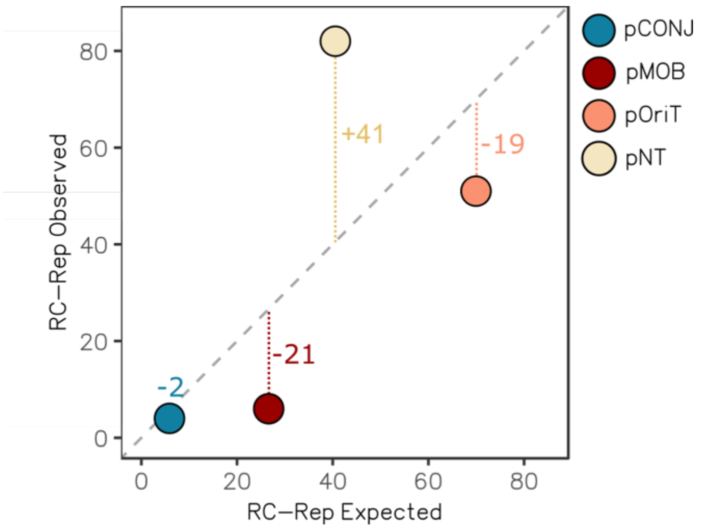


**Supplementary Figure 8. Identification of RC-Rep proteins in *S. aureus*.** Observed number of RC-Rep-encoding plasmids (y-axis) compared to the expected distribution (x-axis). Each dot represents a plasmid type. The numbers next to vertical dashed lines indicate the difference between the observed and the expected.

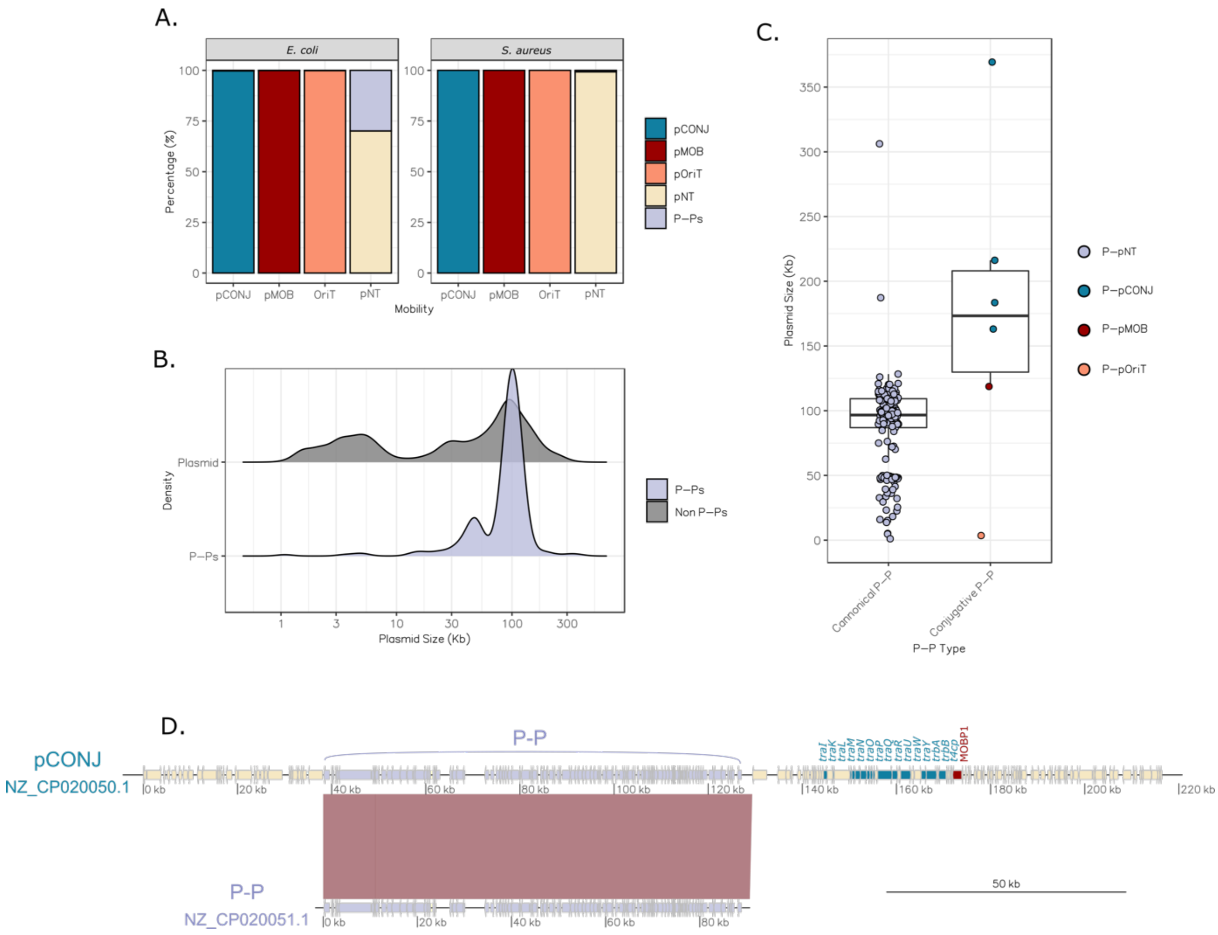


**Supplementary Figure 9. Identification of phage-plasmids (P-Ps). A.** Fraction of P-Ps (purple) identified in *E. coli* plasmids (left facet) and *S. aureus* (right facet). pCONJ: Conjugative plasmid; pMOB: Mobilizable plasmid; pOriT: *oriT*-encoding MOBless plasmid; pNT: putatively non-transmissible plasmids (pMOBless without *oriT*). **B.** Size distribution of P-Ps compared to the size of canonical plasmids. The curves were drawn using a kernel density estimate and scaled to improve the clarity of the representation. **C.** Plasmid size of the typical phage-plasmid (canonical P-P) and the P-Ps identified as pCONJ, pMOB and pOriT. **D.** Alignment of a canonical P-P (no conjugative elements) with a plasmid that is both pCONJ and P-P. The violet arrows indicate the genes associated to the P-P, the blue arrows the MPF genes, and the red arrow the relaxase.

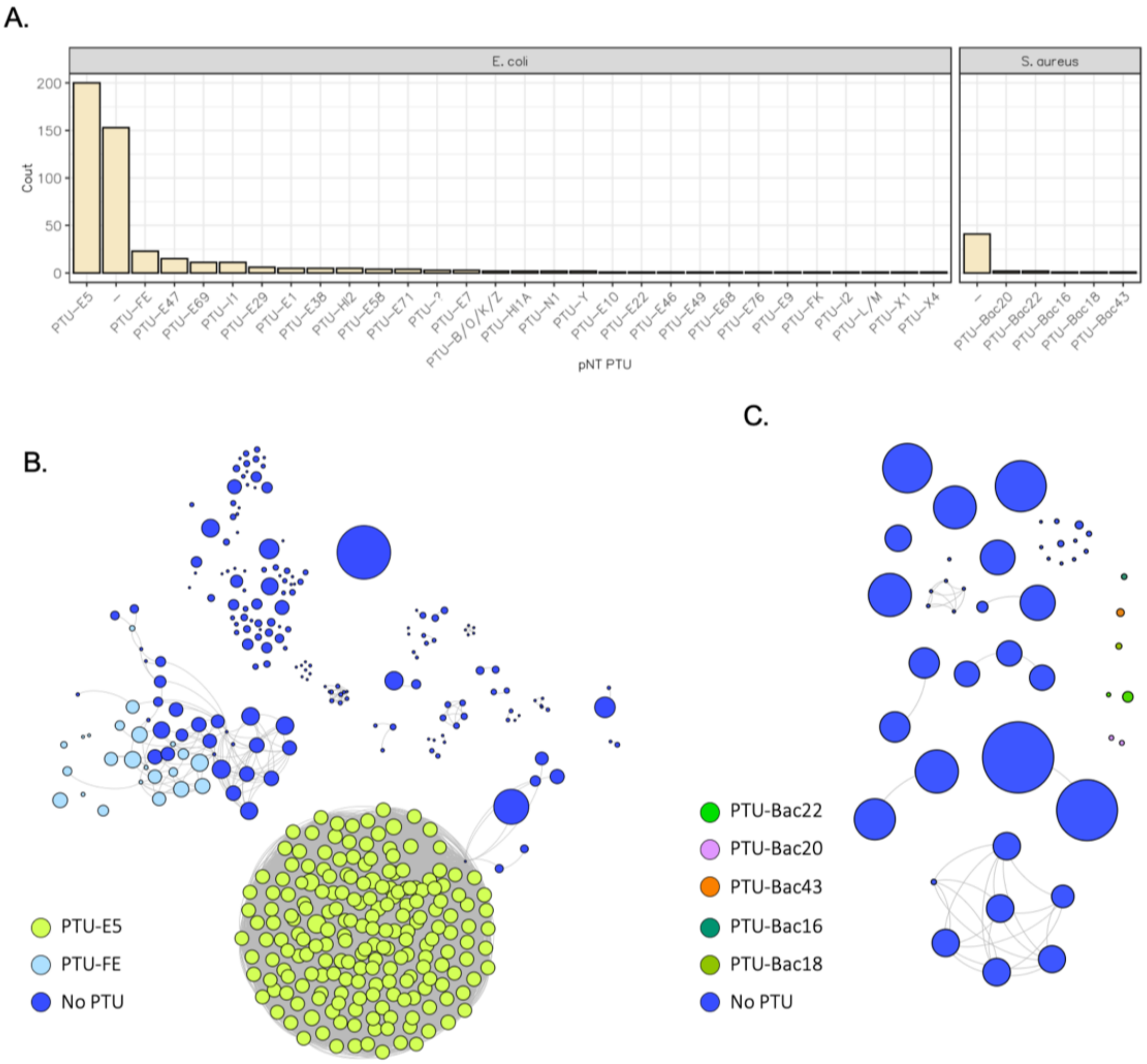


**Supplementary Figure 10. Analysis of putatively non-transmissible plasmids (pNTs). A.** Plasmid Taxonomic Units (PTUs) identified among pNTs. The symbol '-' indicates that no PTU has been assigned. **B.** Graph showing the association between the *E. coli* pNTs assigned to the PTU E5, FE and those without a PTU. The nodes represent the plasmids, colored according to the PTU, whereas the edges indicate a wGRR>0.75. **C.** Graph showing the association between all the *S. aureus* pNTs. The nodes represent the plasmids and the edges wGRRs associations >0.75.

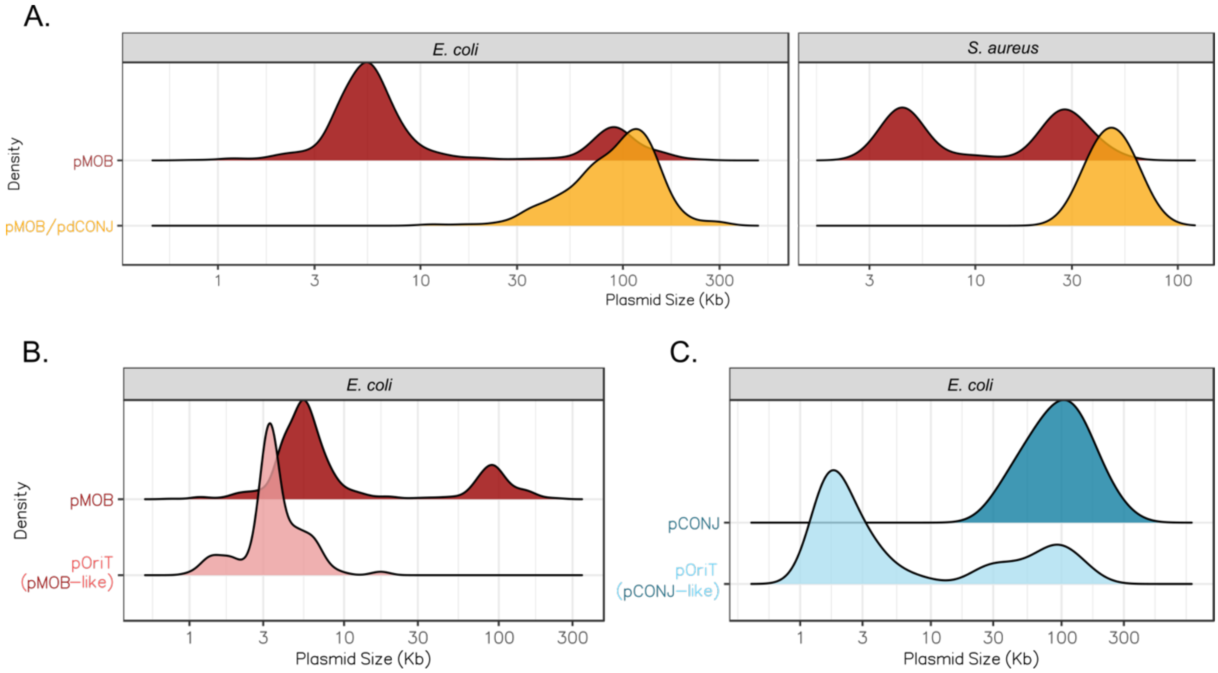


**Supplementary Figure 11. Plasmid size according to the mobility.** The curves were drawn using a scaled kernel density to simplify the representation. **A.** Size distribution of pMOB. pMOB are split into those recently evolved from pCONJ (pMOB/pdCONJ, yellow) or not (pMOB, red). **B.** Size distribution of pMOB (non-pdCONJ) plasmids and pOriTs with pMOB-like *oriTs*. **C.** Size distribution of pCONJ and pOriTs with pCONJ-like *oriTs*. **B.** and **C.** only apply to *E. coli*.

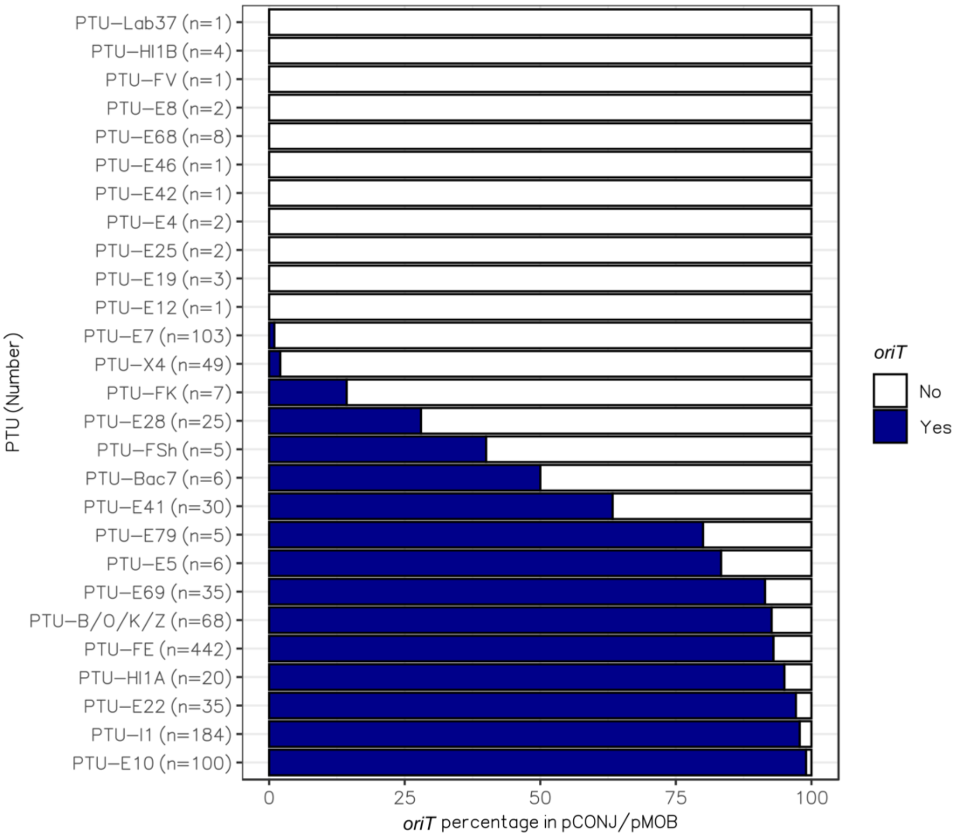


**Supplementary Figure 12. PTUs with non-identified *oriTs*.** Proportion of relaxase-encoding plasmids (pCONJ+pMOB) in each PTU categorized in terms of having an identifiable *oriT* (blue) or not (white). Only PTUs with a at least one non-identified *oriT* are shown. Note that most PTU lacking *oriT*s have very few plasmids (indicated as n=N).

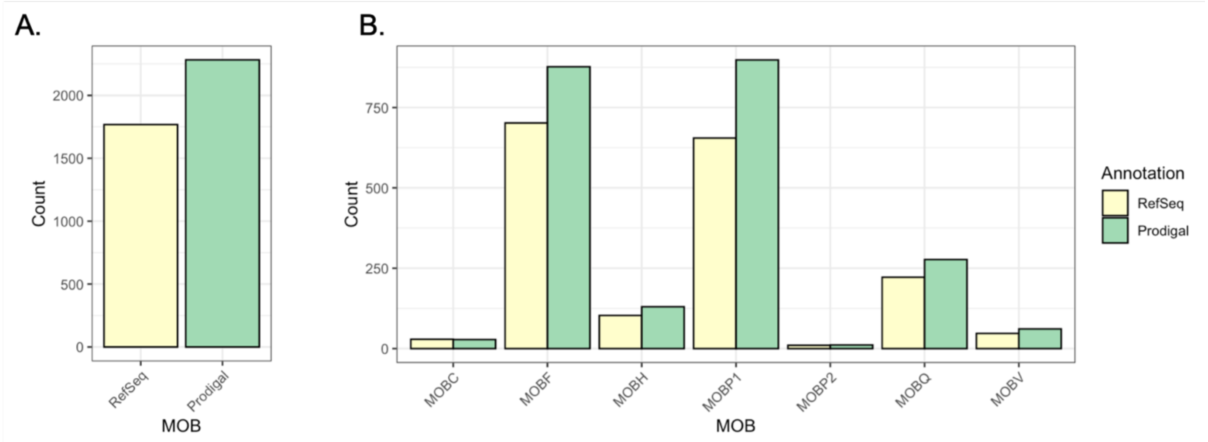


**Supplementary Figure 13. Relaxase identification.** Relaxase identification using the RefSeq annotation retrieved in this study compared to the re-annotation using Prodigal (see Methods). **A.** Overall number of relaxases identified. **B.** Number of relaxases retrieved by the different HMM profiles.

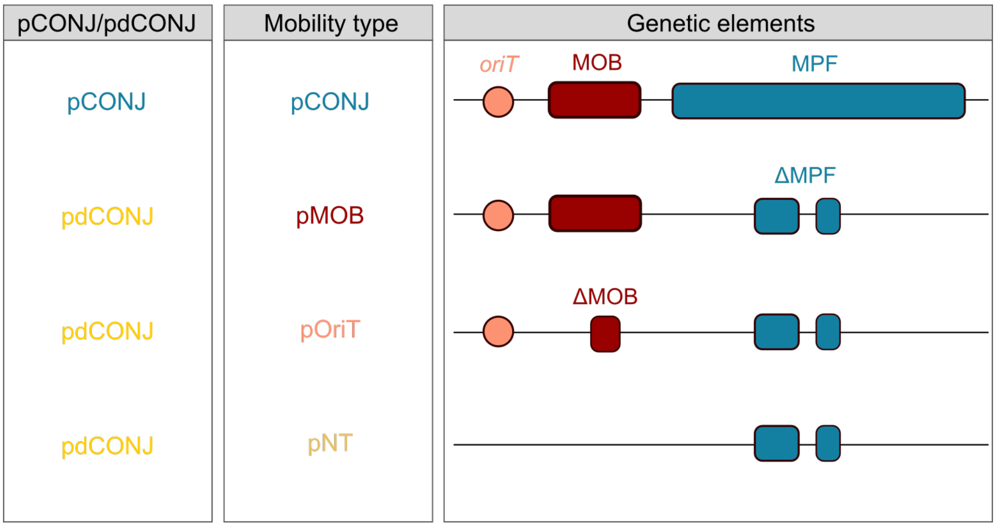


**Supplementary Figure 14. Decayed conjugative plasmids.** Representation of a conjugative plasmid (pCONJ) and different types of decay conjugative plasmids (pdCONJ). The *oriT* is represented with an orange circle, the relaxase with a red rectangle (MOB) and the MPF system with a blue one. ΔMOB and ΔMPF represent incomplete relaxases or MPF systems, respectively. We describe the classification of plasmids in terms of mobility as: conjugative (pCONJ), relaxase-encoding mobilizable (pMOB), *oriT*-bearing mobilizable (pOriT) or putative non-transmissible (pNT).

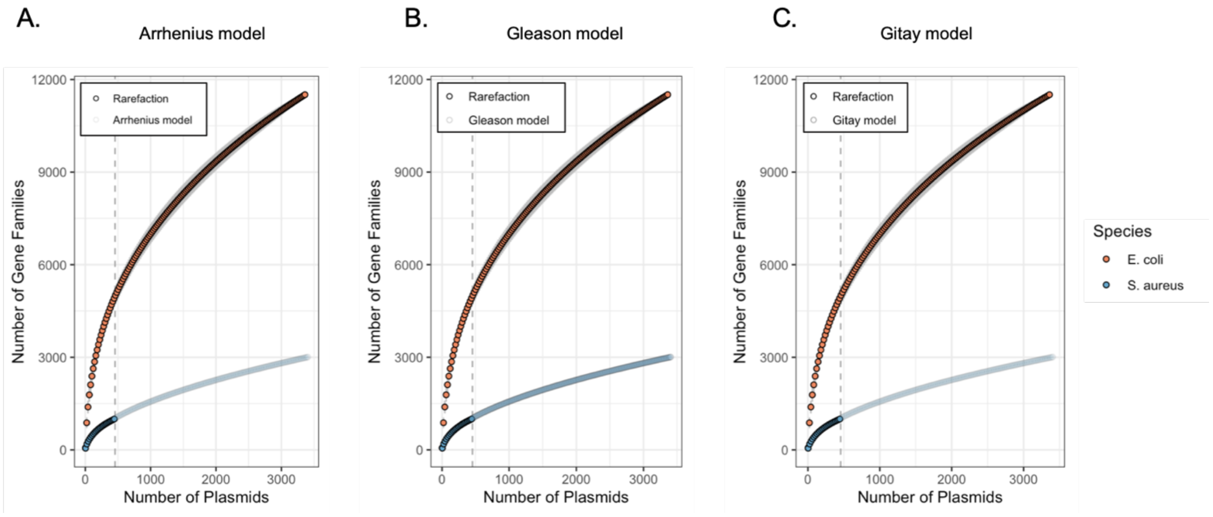


**Supplementary Figure 15. Pangenome analysis of *E. coli* and *S. aureus* plasmids.** The vertical dashed grey line at x = 455 represents the number of plasmids from which *S. aureus* pangenome is inferred following three different non-linear models: **A.** Arrhenius model**, B.** Gleason model, **C.** Gitay model.

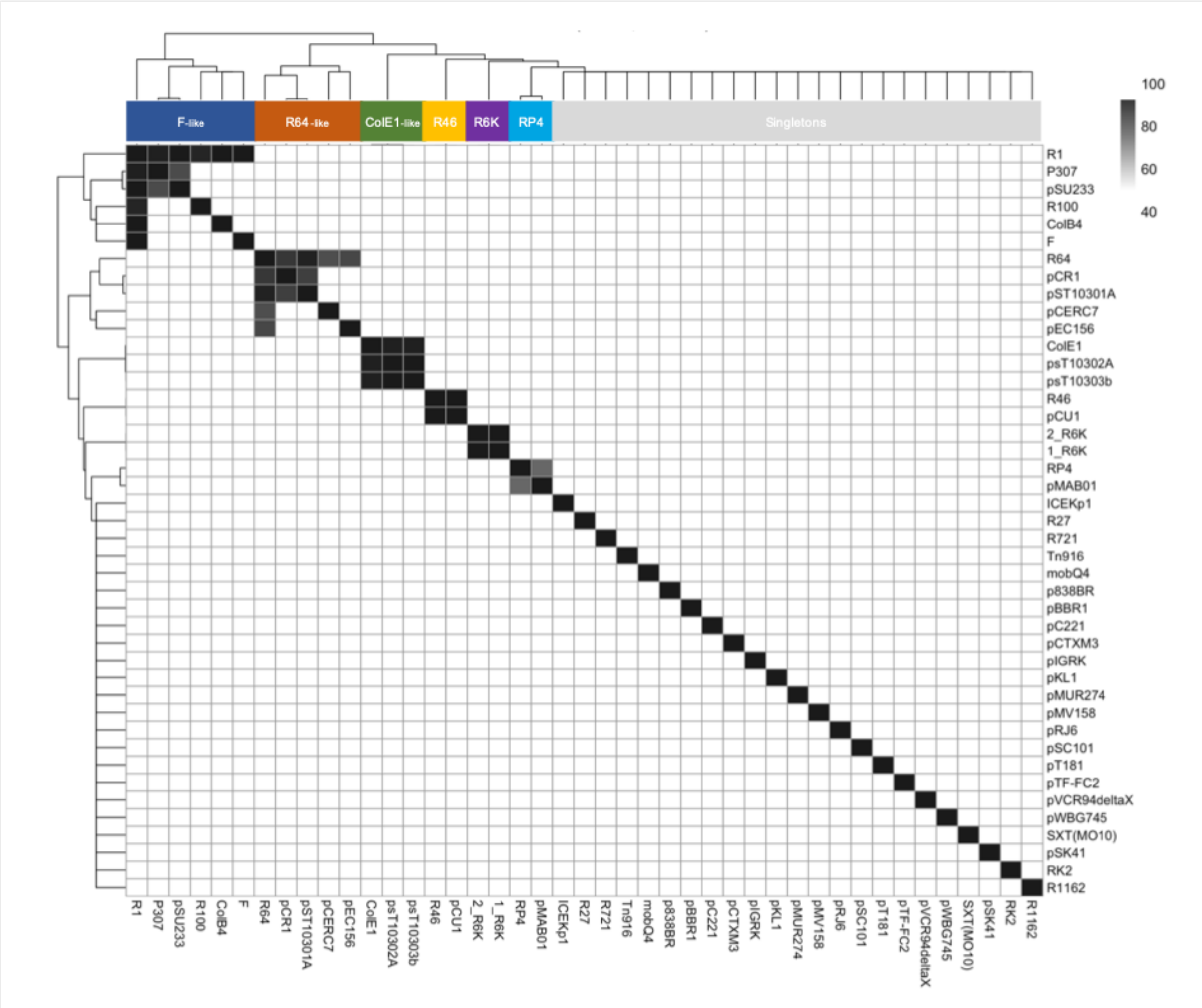


**Supplementary Figure 16. *oriT* families among the origins of transfer identified in the plasmid collection.** The color of the cells indicates the identity (%) of the pairs, being the legend at the right of the figure. Clusters are shown at the top of the heatmap: F-like, R64-like, ColE1-like, R46-like, R6K-like and RP4-like.
